# Preclinical studies with ground germinated barley (GGB) for oral enzyme replacement therapy (Oral-ERT) in Pompe disease knockout mice

**DOI:** 10.1101/2023.12.27.573457

**Authors:** Frank Martiniuk, Adra Mack, Justin Martiniuk, Peter Meinke, Benedikt Schoser, Gongshe Hu, Angelo Kambitsis, John Arvanitopoulos, Elena Arvanitopoulos, Kam-Meng Tchou-Wong

## Abstract

Genetic deficiency of lysosomal acid maltase or acid α-glucosidase (GAA) results in the orphan disease known as glycogen storage disease type II or acid maltase deficiency (AMD) or Pompe disease (PD), encompassing at least four clinical subtypes of varying severity. PD results from mutations in the *GAA* gene and deficient GAA activity, resulting in the accumulation of glycogen in tissues (primarily muscle) and characterized by progressive skeletal muscle weakness and respiratory insufficiency. The current approved enzyme replacement therapy (ERT) for PD is via intravenous infusion of a recombinant human GAA (rhGAA) secreted by CHO cells (Myozyme, Sanofi-Genzyme) given once every 2 weeks and has shown varying efficacy in patients. Although the current ERT has proven to be very efficient in rescuing cardiac abnormalities and extending the life span of infants, the response in skeletal muscle is variable. In late-onset patients, only mild improvements in motor and respiratory functions have been achieved and the current ERT is unsatisfactory in the reversal of skeletal muscle pathology. Additional challenges for ERT include insufficient targeting/uptake of enzyme into disease-relevant tissues, poor tolerability due to severe ERT-mediated anaphylactic and immunologic reactions and the prohibitively high cost of lifelong ERT ($250-500K/year adult patient). A consensus at a Nov.-2019 US Acid Maltase Deficiency Association conference suggested that a multi-pronged approach including gene therapy, diet, exercise, etc. must be evaluated for a successful treatment. Our objective is to develop an innovative and affordable approach via barley GAA (bGAA) from ground germinated barley (**GGB**) or liquid GGB (**L-GGB**) for Oral-ERT for PD or as a daily supplement to Myozyme. To this end, we hypothesize that a bGAA produced in germinated barley can be ingested daily that allows the maintenance of a therapeutic level of enzyme. We have shown in extensive preliminary data that **GGB** or **L-GGB** was (1) enzymatically active, (2) was taken up by GAA KO mice and human WBCs to reverse the enzyme defect that was blocked by mannose-6-phosphate, (3) hydrolyzed glycogen, (4) increased significant changes in the clinical phenotype towards the WT levels in GAA KO mice dose-dependently, (5) taken up by PD myoblasts, lymphoid/fibroblasts cells to reverse the defect, (6) bGAA was ∼70kD, (7) K_m_, V_max_, pH optima, inhibitors and kinetics was similar to human placental GAA and an rhGAA and (8) was strain specific.

## INTRODUCTION

Genetic deficiency of the lysosomal acid maltase or acid α-glucosidase (GAA) results in acid maltase deficiency (AMD) or Pompe disease (PD) encompassing at least four clinical subtypes of varying severity (infantile; childhood, juvenile and adult onset)(**1**). PD exhibits intracellular accumulation of glycogen in multiple tissues with skeletal muscle being the primary target, manifesting as myopathy and cardiomyopathy (**2–5**). The classic infantile form (IOPD) exhibits no enzyme activity, muscle weakness, feeding difficulties and hypertrophic cardiomyopathy, leading to death in the first year (**6–8**). The adult-onset form (LOPD) has partial enzyme deficiency, manifesting with slowly progressive muscle weakness leading to wheelchair/ventilator dependence and premature death from ventilator insufficiency (**9–11**). The incidence of people living with PD is estimated at 1:40,000 (**12**)(Orpha number-ORPHA365 at www.orpha.net). Recently, Park (**13**) re-calculated the carrier frequency (pCF) and predicted genetic prevalence (pGP) using general population databases based on the proportion of causative genotypes. Total pCF and pGP were 1.3% (1 in 77) and 1:23,232. The highest pGP was in the E. Asian population at 1:12,125, followed by Non-Finnish European (1:13,756), Ashkenazi Jewish (1:22,851), African/African-American (1:26,560), Latino/Admixed American (1:57,620), S. Asian (1:93,087) and Finnish (1:1,056,444). Unfortunately, the number of people in each population enrolled may not reflect the real-world population.

ERT with a recombinant human GAA (rhGAA) secreted by CHO cells (alglucosidase alfa/Myozyme/Lumizyme, Genzyme/Sanofi Corp.) infused every two weeks is the first approved therapy. Although it is efficient in rescuing cardiac abnormalities and extending the life span of infants, the response in skeletal muscle is variable (marketed dose of 20 mg/kg/every 2 weeks-iv) via the cation-independent mannose 6-phosphate receptor (**14–17**). In IOPD, muscle pathology and degree of glycogen deposition correlates with the severity of symptoms and the earlier ERT is introduced, the better chance of response (**14**). In LOPD, mild improvements in motor and respiratory functions have been achieved which is unsatisfactory in the reversal of skeletal muscle pathology (**14–17**). In LOPD, the ability for muscle to metabolize extra-lysosomal as well as intra-lysosomal glycogen is impaired. Lysosomal glycogen accumulation leads to multiple secondary abnormalities (autophagy, lipofuscin, mitochondria, trafficking and signaling) that may be amenable by long-term therapy. ERT usually begins when the patients are symptomatic and these secondary problems are already present contributing to inefficient delivery/uptake of alglucosidase alfa to muscle (**18**). The outcome of infantile patients is determined by many factors, among which are age, disease severity at ERT, genotype, genotype dependent CRIM status and the height of the antibody response. High and sustained antibody titers need to be prevented to achieve a good response to ERT, however titers varied substantially between patients and does not strictly correlate with the patients’ CRIM status. Only infantile patients can be either CRIM^neg^ or CRIM**^+^**. Since all juvenile and adult patients make at least some amount of residual GAA, they are always CRIM**^+^**. About 40% of infantile patients are CRIM^neg^ while the remaining 60% are CRIM**^+^**. The immune response may be minimized by early start of ERT and pretreatment by an immune tolerance induction regime using e.g. a combination of rituximab, methotrexate, bortezomib and IV immunoglobulins. Up to 20% of adult patients develop high titers on ERT, 40% intermediate and 40% none or low titers (**19–40**). In August 2021, the FDA approved Nexviazyme (avalglucosidase alfa-ngpt) for the biweekly treatment of patients >1yr and older. Nexviazyme has ∼15x mannose-6-phosphate (M6P) compared to alglucosidase alfa. Nexviazyme in trials with LOPD, improvements in respiratory function and walking distance, but not statistical superiority of Nexviazyme (**41–44**). Cohen et al. (**45**) recently performed a Phase 1 clinical trial of *in utero* ERT with Myozyme for a single IOPD (CRIM^neg^) receiving six *in utero* ERT starting at 24 weeks of gestation via umbilical vein followed by standard postnatal ERT. The newborn patient had normal cardiac and motor function, was meeting developmental milestones, had normal biomarker levels, decreased placental glycogen deposits, was feeding and growing normally at 13 months (**45,46**), had low levels of antidrug antibodies at week 28 of gestation, but postnatal ERT development of high antidrug antibody titers which was managed with inhibitors (**18,45–46**).

In adults, antibody formation does not interfere with rhGAA efficacy in the majority of patients, however may be associated with infusion-associated reactions (IARs) and may be attenuated by the IVS1/delex18 (c.2481+102_2646+31del, expression of a truncated protein) GAA genotype (**37**). Some patients experienced IARs due to bronchial spasm with flushing and pulmonary artery blockage during infusion. Several studies suggest that unchanged endogenous GAA protein expression lowers the chance of forming antibodies to recombinant proteins. Gutschmidt et al. (**47**) found that ERT in LOPD clinical outcomes, particularly lung function, muscle strength and walking capability tend to deteriorate over time, indicating that additional efforts must be made to improve ERT effectiveness and supplement therapies developed. Korlimarla et al. (**48**) used a standardized behavior checklists as screening tools for the early identification and treatment of behavior, emotional and social concerns in children with PD. Although alglucosidase alfa has been a wonderful first step in treating people with PD, it has revealed subtle aspects that must be dealt with for successful treatment (**18,50–54**). ERT has unmasked previously unrecognized clinical manifestations include tracheo-bronchomalacia, vascular aneurysms and GI discomfort that impacts smooth muscle. Persistent smooth muscle pathology has a substantial impact on quality of life and leads to life threatening complications. In addition to airway smooth muscle weakness, vascular deterioration, GI discomfort and loss of genitourinary control have been observed. Cerebral and aortic aneurysms have caused microhemorrhages leading to symptoms ranging from headaches and numbness to coma and death (**55–61**). PD results in subsequent pathology in smooth muscle cells may lead to life-threatening complications if not properly treated (**62**). In GAA KO mice, there is increased glycogen in smooth muscle cells of the aorta, trachea, esophagus, stomach and bladder plus increases in lysosome membrane protein (LAMP1) and autophagosome membrane protein (LC3). More importantly, lifelong treatment ($250-500k/year/adult) can be prohibitively expensive resulting in the reluctance of insurance companies to reimburse costs for adult patients (**63**) underlining the demand for more economical production and novel delivery strategies for treatment. A consensus at a 2019 US Acid Maltase Deficiency Association conference in San Antonio, TX, suggested that a multi-pronged approach including gene therapy, diet, exercise, etc. must be evaluated for a successful treatment of PD.

Previously, we expressed the *hGAA* in tobacco seeds (tobrhGAA)(**64,65**) as a plant made pharmaceutical that was enzymatically active. In PD KO mice, daily Oral-ERT with ground tobrhGAA seeds showed significant reversal of fore-limb and hind-limb muscle weakness, increased motor coordination/balance/strength and mobility, improved spontaneous learning, increased GAA activity in tissues, reduced glycogen in tissues and negligible antibodies to hGAA. The tobrhGAA was taken up in PD fibroblast, lymphoid and myoblast cells to reverse the enzymatic defect. Enzyme kinetics compared favorably or superior to placental hGAA, plus alglucosidase alfa or other rhGAAs for K_m_, V_max_, pH optima, thermal heat stability and IC_50_ for inhibitors. The tobrhGAA in seeds was extremely stable stored for 15 years at room temperature. Thus, Oral-ERT with ground tobrhGAA seeds is an innovative approach that overcomes some of the challenges of alglucosidase alfa-ERT and provides a more effective, safe and significantly less expensive treatment (**65**). Current ERT for PD worldwide especially in areas where access to a clinical setting for the biweekly IV administration of Myozyme is lacking. We estimate that an adult PD patients will need to take orally 3-4 grams daily of tobrhGAA at a cost of ∼1-3% of Myozyme depending on how it is branded and marketed to maintain a sustained GAA level of 5-10% of normal. This product is considered a drug as a GMO dietary/nutriceutical supplement by the FDA and must undergo clinical trials, etc for use in humans unless we can get it reclassified and this obstacle is removed. A retrospective study indicates health care costs for rare diseases are >4x for patients without a rare disease (**65a**). About 10% of rare diseases have an FDA approved therapy and underscore an urgent need for more research and earlier, more accurate diagnoses of and interventions for rare diseases. Most of the ∼7,000 to 10,000 known rare diseases disproportionately affect children, adolescents and young adults. Each year, ∼250 rare genetic diseases are added. One in 135 babies are born with a genetic condition that needs to be treated in childhood (**65b**). Owen et al. (**65c**) in a cohort study, WGS identified single-locus genetic diseases in a high proportion (41%) of 112 infants who died between 2015 and 2020 while genetic diseases accounted for 28% among all 311 infant deaths without WGS. Individually, most rare diseases might affect only a few hundred to a few thousand worldwide. However, collectively rare diseases are common, affecting an estimated 25 million to 30 million patients in the USA. Many of these diseases are genetic, life-threatening and hard to diagnose and treat. ERT is available for 28 disorders. In the USA, costs ranged from $9,000-$140,000/year/patient with a total yearly medical costs of ∼$400 billion of all rare disease patients. A new FDA legislation removes the mandate of drugs to be tested on animals before moving onto human clinical trials that will save thousands of animals from unnecessary suffering plus will make drug screening faster and more efficient (www.npr.org/2023/01/12/1148529799/fda-animal-testing-pharmaceuticals-drugdevelopment https://www.paul.senate.gov/news/dr-pauls-bipartisan-fda-modernization-act-20-end-animal-testing-mandates-included-2022-year).

Seed dormancy is a developmental checkpoint that prevents mature seeds from germinating under conditions that are otherwise favorable for germination. Temperature and light are the most relevant environmental factors that regulate seed dormancy and germination. These environmental cues can trigger molecular and physiological responses including hormone signaling, particularly that of abscisic acid and gibberellin. The balance between the content and sensitivity of these hormones is the key to the regulation of seed dormancy (**66**). Gibberellic acid (GA) is a simple gibberellin, breaking seed dormancy, promoting seed maturation, growth and elongation of cells. GA stimulates the cells of germinating seeds to produce mRNA molecules and post-translational events that code for hydrolytic enzymes. It is usually used in concentrations between 0.01 and 10mg/l (**67**). Yu and Wang (**68**) review advances in the molecular connections between miRNAs and GA with an emphasis on the two miRNAs, miR156 and miR159. Briggs et al. (**69**) and Vieira et al. (**70**) demonstrated that treatment of barley endosperm with GA stimulates the production of many hydrolytic enzymes and the release of sugars, polypeptides and inorganic phosphate into the culture medium with the endosperm of the treated preparations frequently disaggregating completely. Chen et al. (**71**) showed that GA at 0.1DM stimulates amylase synthesis in dormant *Avena fatua* seeds without inducing germination in early hours (0-14 hours). Briggs (**72**) and Riley et al. (**73**) reviewed uses of GA to breaking barley dormancy, increasing malt extract and reducing malting time in the aleurone layer. Tibbot et al. (**74**) produced expression of enzymatically active recombinant bGAA in yeast and immunological detection of α-glucosidase from seed tissue. An α-glucosidase cDNA clone derived from barley aleurone tissue was expressed in *P. pastoris* and *E. coli*. From yeast, the enzyme hydrolyzed maltose over the pH range 3.5–6.3 with an optimum at pH 4.3, classifying the enzyme as a GAA with a 81kD GAA prominent accompanied by a 14-fold increase in bGAA. The 13 amino acid catalytic region from spinach and sugar beet α-glucosidases are identical to bGAA and hGAA (DGX**<u>WIDMNE</u>**XSNF)(**74**). Frandsen et al. (**75**) purified a high-isoelectric-point (pI) α-glucosidase from barley (*Hordeum vulgare*) malt. The enzyme had high activity toward maltose (kcat = 25 s^-1^), with an optimum at pH 4.5 and catalyzed the hydrolysis. Acarbose was a strong inhibitor (Ki = 1.5 M). Mass spectrometry assigned the 92kD protein to a barley cDNA (GenBank accession no. U22450) that appears to encode a GAA (**75**). Muslin et al. (**76**) overexpressed, purified and characterized a bGAA secreted by *Pichia pastoris.* GAA (EC 3.2.1.20) are recognized as important in starch degradation during cereal seed germination. bGAA (*Hordeum vulgare*) expressed in *Pichia pastoris* had a pH optima for maltose at 4 (**76**). Andriotis et al. (**77**) recently showed that the high and low pI bGAAs may be encoded by only one gene with the two isoforms are due to proteolysis and post-translational modifications. Maltase is the enzyme that is responsible for metabolizing starches into glucose in the intestinal lining of the stomach (**78**).

Our objective is to develop an innovative and affordable approach via bGAA from ground germinated barley (**GGB**) or liquid GGB (**L-GGB**) for Oral-ERT for PD or as a daily supplement to Myozyme. To this end, we hypothesize that a bGAA produced in germinated barley can be ingested daily that allows the maintenance of a therapeutic level of enzyme. We have shown in extensive data that **GGB** or **L-GGB** was (1) enzymatically active, (2) was taken up by GAA KO and human WBCs to reverse the enzyme defect that was blocked by mannose-6-phosphate, (3) hydrolyzed glycogen, (4) increased significant changes in the clinical phenotype towards the WT levels in GAA KO mice dose-dependently, (5) taken up by PD myoblast, lymphoid and fibroblast cells to reverse the defect, (6) was ∼70kD, (7) K_m_, V_max_, pH optima, inhibitors and kinetics was similar to human placental GAA and rhGAA and (8) was strain specific.

## MATERIAL and METHODS

### Germination and preparation of barley

Barley (*Hordeum vulgare L*.); strain Morex or Robust are surface sterilized by soaking in 5% bleach for 5 minutes, washed with water and placed on paper towels saturated with GA (0.05mg/ml; 0.14mM; 50ppm) for 1 day at RT, replace paper towels with fresh water followed by germination for 7 days in the dark at RT (**67–78**). Germinated seeds have the shoots/roots removed and stored at −20°C. We processed 8-day wet seeds by drying in a food-dryer, ground until a fine powder, UV sterilized in a germicidal incubator for 30 minutes and stored at RT = **GGB**. **L-GGB** was prepared by suspension of **GGB** or shoot/rootless geerminated seeds in 0.001 M sodium phosphate pH 7.5 at 1.2 volume/weight in a blender. The homogenate is filtered through 2 layers of fine cheesecloth, followed by passing through a 200 mesh nylon membrane and stored at 4°C.

#### Simulated Stomach and Small Intestinal Environments

To mimic the stomach and small intestine (SI) environment, we exposed the 100 mg of whole or milled seeds to physiologic conditions/times to pepsin (stomach) followed by trypsin/chymotrypsin (small intestine)(**79,80**). Samples were added to 300 μg/ml pepsin at pH 4.0 for 60 min., adjusted pH to 6.5 and then add trypsin at 800 μg/ml and chymotrypsin at 700 μg/ml for 60 min at 37°C followed by GAA assay.

#### Enzyme Assay

Ground seeds or tissues were resuspended in 10 mM sodium phosphate, pH 7.5, frozen and thawed 3x and centrifuged at 13,000g for 10 min. The supernatants were assayed for GAA using the artificial substrate 4-methylumbelliferyl-α-D-glucoside (4-MU-Glc, 1 mg/ml) at pH 4.0 (0.5 M sodium acetate) and as an internal control, neutral alpha glucosidase (NAG) at pH 7.5 (0.5 M sodium phosphate) for 18 hours. Fluorescence was determined in a fluorometer (excitation-360_nm_ and emission-460_nm_)(Sequoia-Turner) as previously described (**81**).

#### Biochemical/Enzyme Kinetic Analyses

We compared a lysate of seeds to a rhGAA (R&D Systems) and mature placental human GAA for specific activity using 4-MU-Glc at pH 4.0, maltose and glycogen, pH optima, inhibitors (acarbose, castanospermine, deoxynojirimycin, miglitol, voglibose) and heat stability (**82–94**) using standard enzyme kinetic methods. (http://www.ic50.tk/kmvmax.html, http://www.ic50.tk/kmvmax.html, https://www.mdapp.co/michaelis-menten-equation-calculator-431/, http://calistry.org/calculate/michaelis-menten-Equation, http://www.ic50.tk/)(82–97). The K_m_, V_max_ and IC_50_ for the 50-100kD fraction for 4-MU-Glc, glycogen, maltose and 4-MU-Glc/glycogen using software-(http://www.ic50.tk/kmvmax.html, http://www.ic50.tk/kmvmax.html, https://www.mdapp.co/michaelis-menten-equation-calculator-431/, http://calistry.org/calculate/michaelis-menten-Equation, http://www.ic50.tk/)(82–97).

### Uptake of L-GGB by WBCs from Human and GAA KO Mice

**L-GGB** uptake by white blood cells from a human and GAA KO mice was tested ± 5mM mannose-6 phosphate. Mannose-6 phosphate treatment blocks uptake by the M6P receptor (**98–100**). hGAA was used as a positive control. Anti-coagulate-EDTA was added at 1.5 to 2.0 mg/ml of whole blood. Human and mouse KO blood (0.15ml) was mixed with a 14x RBC lysing solution (NH_4_Cl (ammonium chloride)-8.02g; NaHCO_3_ (sodium bicarbonate)-0.84g; EDTA (disodium)-0.37g to 100ml with water) in a 1.5 ml microfuge tube, inverted for 10 minutes at RT until clear red followed by spinning for 1 minute in a microfuge. The supernatant was decanted, cells were washed with phosphate buffered saline (PBS) and re-suspend in 2.4 ml complete DMEM-10%FBS (Life Technologies). Cells were divided into 6 sterile microfuge tubes, 70 μl of **L-GGB** or 70 μl placental hGAA or 70 μl PBS was added and placed at 37°C for 18 hours. In one set, M6P (5mM) was added to block uptake. Lysed cells were assayed for GAA and NAG.

#### Uptake of tobrhGAA by PD Human Myoblast, Fibroblast and Lymphoid Cell Lines

Human lymphoid (GM6314, GM13793, GM14450) or fibroblast (GM4912, GM1935, GM3329) cell lines from infantile or adult PD were maintained in 15% fetal bovine serum, RPMI 1640 or DMEM supplemented with glutamine, penicillin and streptomycin at 37°C-5% CO_2_. Cells were plated at 0.3-0.4 x 10^6^/6-well in 1.5 ml media-24 hours before addition of varying amounts of **L-GGB** or others GAA formulations (**101**). Cells were harvested after various hours of exposure, washed with PBS, lysed by addition of 0.5 ml of 0.01 M sodium phosphate pH 7.5, frozen and thawed 3x, spun for 5 minutes to clarify and assayed for human GAA and NAG as described above. A human PD myoblast cell line (homozygous for the IVS1 c.-32–13t>g)(**102**) and normal skeletal muscle cells were plated at 0.3 to 0.4 x 10^6^ in a 6-well plate with PromoCell skeletal muscle growth media. Cells were exposed to **L-GGB** for 48 hour and assayed. Mock treated GAA and normal myoblast cells were controls plus cells treated with equivalent amounts of a rhGAA (R&D Systems #8329-GH-025). Simultaneously, we measured cell proliferation with the Cayman MTT cell proliferation assay (#10009365).

#### Studies GAA KO Mice

We utilized the GAA KO mouse with the exon 6^neo^ disruption (**103**), wild-type BALB/c or 129/J or GAA KO mice mock-treated with PBS. GAA KO mice (∼4-6 months old) were given oral administrations every other day of **L-GGB** (∼0.1μg/g bGAA) mixed 3:1 with apple juice as a safe, non-invasive oral administration technique (**104**), grip-strength measured by a grip-strength meter (GSM)(Columbus Inst. OH) and vertical hang-time (Raben et al., **103**). Mice were sacrificed at different times and tissues assayed for GAA and NAG and compared to wild-type mice and mock (PBS) treated GAA KO mice. Additionally, GAA KO mice were fed either 50 mg or 150 mg **GGB** daily mixed with peanut butter in petri dishes (12:12 L/D photoperiod)(**64**)(age ∼5 months)(B6,129-Gaa^tm1Rabn^/J mice, **103**). Tissues, urine, blood slides, weight and serum were collected. Mice were tested every 3 weeks for: motor activity by running wheel (RW-Mini-Mitter Co., Inc., OR); fore-limb muscle strength by grip-strength meter (GSM)(Columbus Inst. OH), motor coordination/balance with a Rotarod (AJATPA Expert Industries, India), open-field mobility by a 5 minute video for distance traveled and spontaneous alternative learning with a T-maze (Stoelting, IL).

#### Assay for Glycogen Content

In a 96-well microtiter plate, add 25 μl sample, 25 μl 0.05 M sodium phosphate pH 6.5, ±0.5 μl amyloglucosidase (Sigma #A-1602), placed overnight at 37°C (**105**). Glycogen standards (rabbit liver, 2 mg/ml) and D-glucose standards were at 400 μM, 200 μM and 100 μM. Add 20 μl of Eton Bioscience glucose assay solution (#1200031002) and incubate for 30 minutes at 37**°**C. The reaction was stopped with 25 μl of 0.5 M acetic acid and read at A_490_.

#### CBC differential

CBC differentials were perform on blood smears on slides with Giemsa-Wright staining.

#### Statistical Analysis

Student t-test for probability associated with a population and standard deviation based upon the population were determined using Microsoft Excel software and considered significant at p = ≤ 0.05.

## RESULTS

### A-Miscellaneous studies

#### A1-Germination and bGAA expression

Barley (*Hordeum vulgare L*.) strain Morex or Robust are surface sterilized by soaking in 5% bleach for 5 minutes, washed with water and placed on paper towels saturated with GA (0.05mg/ml; 0.14mM; 50ppm) for 1 day at RT, replace paper towels with fresh water followed by germination for 7 days in the dark at RT (**67–78**). Germinated seeds have the shoots/roots removed and stored at −20°C. We processed 8-day wet seeds by drying in a food-dryer, ground, UV sterilized and stored at RT. Processed seeds lost ∼75% water weight and no loss in bGAA activity. Thus, 6-wet seeds = 0.5g or 0.15g **GGB**. bGAA and NAG activities were determined using the fluorescent substrate 4-MU-Glc at pH 4.0 for bGAA and pH 7.5 for NAG. bGAA had 0.1μg/g of **GGB**. GA stimulation and a time-course of expression for bGAA activity from Morex and Robust strains showed a maximum activity of bGAA at days 7-8 (**Figure 1**). Un-germinated and generic barley did not show GA stimulation/time-course. **GGB** also hydrolyzed glycogen (K_m_-1.06mg/ml; V_max_-52.7nmole/min/ml) similarly to placental human GAA (**Table 1**).

**Figure 1.**
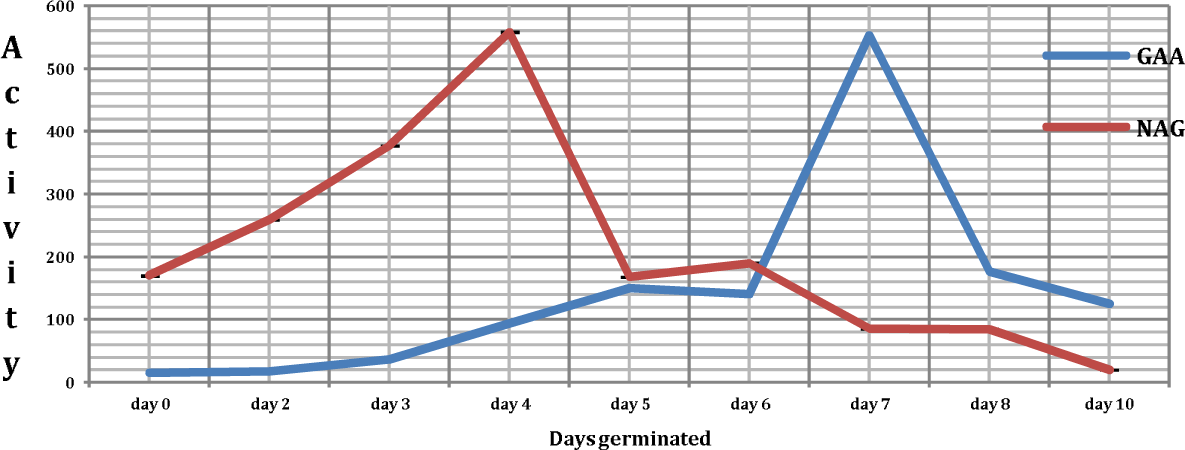
GAA & NAG activity in gibberelin germinated barley is a graph showing GAA and NAG activity in gibberellin germinated barley. Barley GAA (bGAA) activity peaked at days 7-8 of germination in Morex and Robust barley varieties.

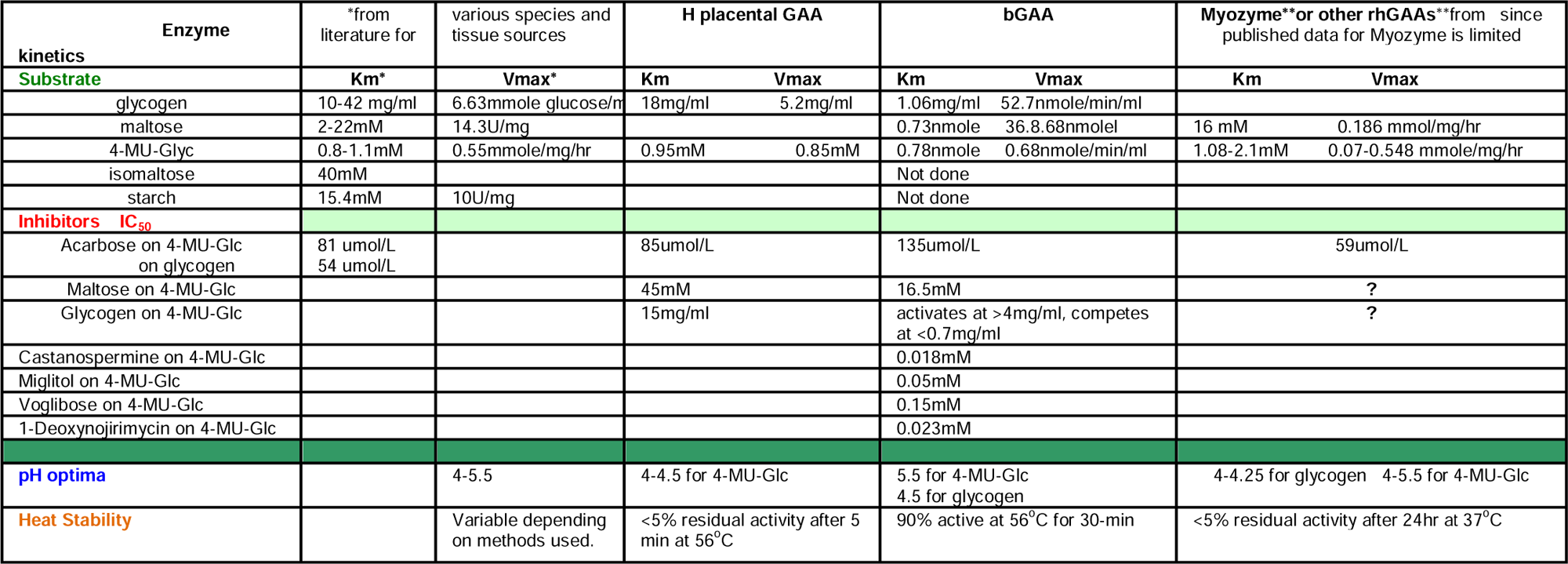
Table 1.

#### A2-GA stimulation and time-course in others grains

We tested 10 other grains (corn, millet, rye, wheat, sorghum, buckwheat, quinoa, rice, sunflower and oats) for GA stimulation and time-course (to day 10) for expression of grain specific GAA to compare to the Morex or Robust strain profiles for bGAA. Assays for GAA and NAG in lysates from these grains for 10-days of germination demonstrated that these grains did not have any substantial increase in GAA or NAG profiles as compared to the Morex and Robust strains. Thus, the bGAA/NAG profile is unique to these barley strains.

#### A3-Long-term stability

**Figure 2** shows that dried vs. non-dried seeds at day 7 germination stored up to 2 months at RT with no reduction in bGAA activity (mean±SD), thus bGAA in germinated seeds is stable at RT for months. Further testing of **GGB** stored for 1.0 years at RT vs. freshly produced that there was less than 5% difference in bGAA activity between the two preparations indicating extreme stability (data not shown). We also found that there was no loss in bGAA activity stored at 4°C for over 1.5 years while bGAA at RT had no loss in activity at 4 months followed by a steady decline to 25% by 12 months (**Figure 3**).

**Figure 2.**
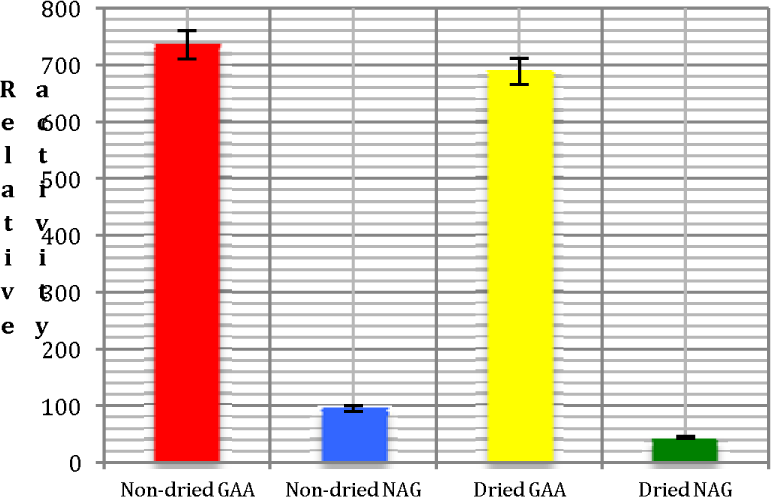
Barley gibberelin germinated and assayed at day 7; non-dried vs. dried seeds stored at RT for 2 months is a bar graph showing barley germinated with gibberellin and assayed at day 7 of germination in non-dried versus dried seeds that were stored for 2 months at room temperature. Analysis of bGAA and NAG activity in the samples showed that there was no reduction in bGAA activity (mean±SD) due to drying and storage.

**Figure 3.**
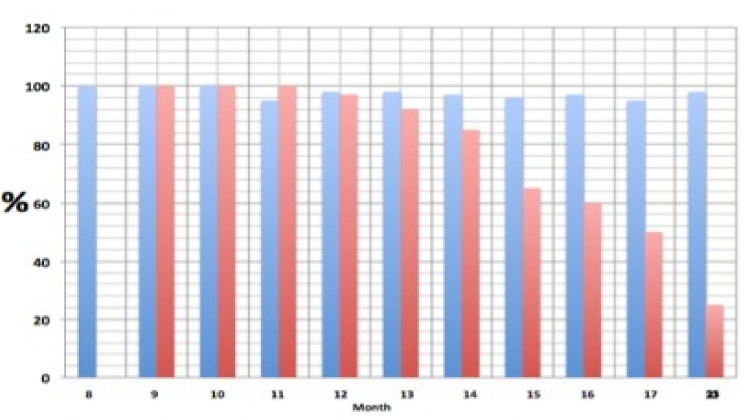
Stability of bGAA Stored at 4oC vs RT is a bar graph showing there was no loss in bGAA activity stored at 4°C for over 1.5 years while bGAA at RT had no loss in activity at 4 months followed by a steady decline to 25% by 12 months. a bar graph showing the heat stability of bGAA. bGAA was treated at three temperatures (56°C, 60°C, or 65°C) for 5, 10, 15, 20, or 30 minutes and assayed for GAA activity. Activity is shown compared to untreated bGAA (100% activity).

#### A4-Determination of the active molecular weight

Partial purification of bGAA from **L-GGB** was performed by size exclusion using Amicon YM 50kD and 100kD membranes (MilliporeSigma, Burlington MA). Chromatography on Sephacryl S200 HR showed bGAA activity eluted at a MW of ∼70kD (**Figure 4**). Standards used for size determination were bovine serum albumin (BSA)(69kD), ovalbumin (OA)(48kD), and cytochrome C (Cyto-C)(14kD).

**Figure 4.**
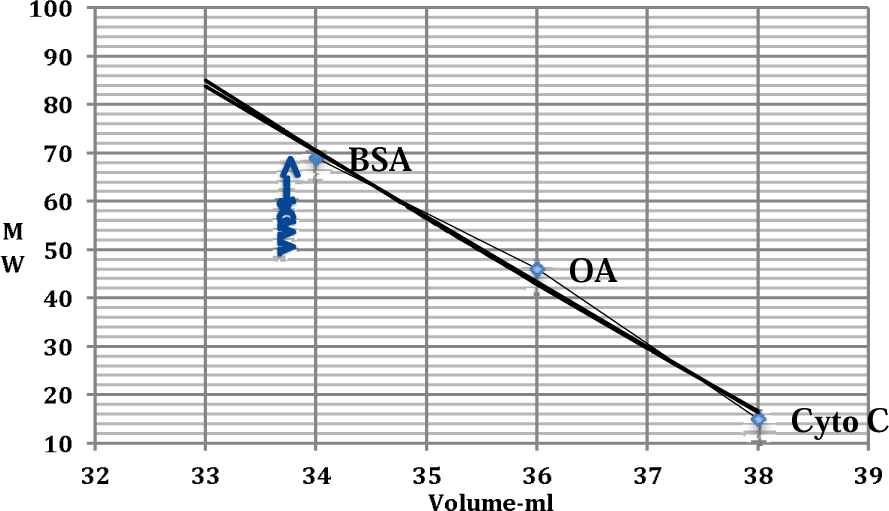
Sephacryl S200-HR column is a bar graph showing chromatography on Sephacryl S200 HR of bGAA activity eluted at a MW of ∼70kD. Standards used for size determination were bovine serum albumin (BSA) (69kD), ovalbumin (OA) (48kD), and cytochrome C (Cyto-C)(14kD). line graph showing determination of bGAA molecular weight by chromatography on Sephacryl S200 HR with an extract of GGB. Peak bGAA activity eluted at a MW of ∼70kD. The bGAA was partially purified by size inclusion between Amicon YM 50kD and 100kD membranes. Standards used for size determination were bovine serum albumin (BSA)(69kD), ovalbumin (OA)(48kD) and cytochrome C (Cyto-C)(14kD).

#### A5-Molecular structure and homology

We compared the *Homo sapiens GAA* mRNA NM_000152.3 of 3782bp and 952 residues vs barley (*Hordeum vulgare*) high pI GAA (AGL97), GenBank:AF118226.1/U22450.1/ AAB02985 for a mRNA of 2738 bp and 877 amino acids. NCBI/BLAST/blastp suite showed a 433/783 (55%) nucleotide homology with both genes containing the putative catalytic seq of **WIDMNE**. NetNGlyc predicted 6 Asn-X-Ser/Thr sequons for N-glycosylated in bGAA at amino acids 191,298,338, 391,471 and 570. hGAA has 7 sites at 140, 233, 390, 470, 652, 882 and 925.

#### A6-Heat stability

Heat stability of purified bGAA was determined by incubation of 25 μl of the 50-100kD bGAA fraction for various times over three temperatures. The temperatures tested were either 56°C or 60°C or 65°C for 5, 10, 15, 20 or 30 minutes at each temperature and assayed for GAA activity. These data show that bGAA is extremely heat stable at 56°C for up to 30 minutes. Stability at 60°C and 65°C declined rapidly over the 30 minute treatment (**Figure 5**).

**Figure 5.**
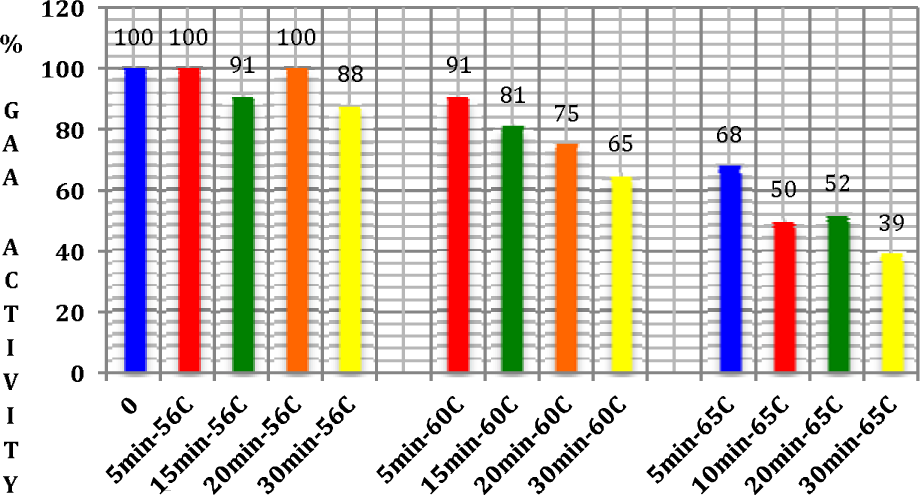
% GAA Heat Stability for Robust Barley 100-50 kD Fraction is a bar graph showing heat stability of purified bGAA by incubation of the 50-100kD bGAA fraction for various times over three temperatures. The temperatures tested were either 56°C or 60°C or 65°C for 5, 10, 15, 20 or 30 minutes at each temperature and assayed for GAA activity.

#### A7-Analysis for biologic contaminates

**L-GGB** were analyzed for biologic contaminates using R-Cards from Roth Bioscience LLC (Goshen, IN, rothbioscience.com) for *Salmonella* species, *E. coli*, yeast/mold and *Listeria* species. Briefly, for specific R-cards, the top plate was lifted, 1 ml of sample was placed on the center of the card, then the card was slowly re-covered with the top plate. After standing for 1 minute so that the the sample was evenly spread, the R-cards were incubated at 37°C overnight for all organisms except yeast which was placed at 25°C for 24-72 hours. Sensitivity was one organism/ml. No biologic contaminates were found in **L-GGB**.

#### A8-Biochemical/enzyme kinetic analyses

We compared the enzyme kinetics for bGAA vs placental hGAA, and an rhGAA (R&D Systems), plus limited data for alfa or other rhGAAs (**86–94**) for K_m_, V_max_, pH optima, thermal heat stability and IC_50_ for inhibitors (acarbose, castanospermine, deoxynojirimycin, miglitol, voglibose). Data showed that showed that bGAA is comparable to placental GAA and superior to alglucosidase alfa and other rhGAAs (**Table 1**, **Figures 6-10**)(http://www.ic50.tk/kmvmax.html, http://www.ic50.tk/kmvmax.html, https://www.mdapp.co/michaelis-menten-equation-calculator-431/, http://calistry.org/calculate/michaelis-menten-Equation, http://www.ic50.tk/)(82–97). **Figures 6-10** show the K_m_, V_max_ and IC_50_ for the 50-100kD fraction for 4-MU-Glc, glycogen, maltose and 4-MU-Glc/glycogen using software-http://calistry.org/calculate/michaelis-menten-Equation, http://www.ic50.tk/) and pH optima for 4-MU-Glc (pH 5.5) and glycogen (pH 4.5).

**Figure 6.**
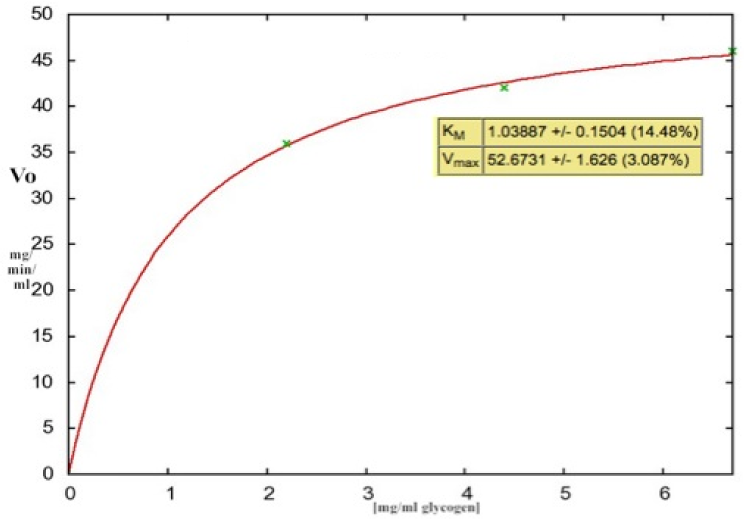
Km & Vmax of glycogen for barley >50kD are bar graphs showing the Km, Vmax and IC50 for the 50-100kD fraction for 4-MU-Glc, glycogen, maltose and 4-MU-Glc/glycogen.

**Figure 7.**
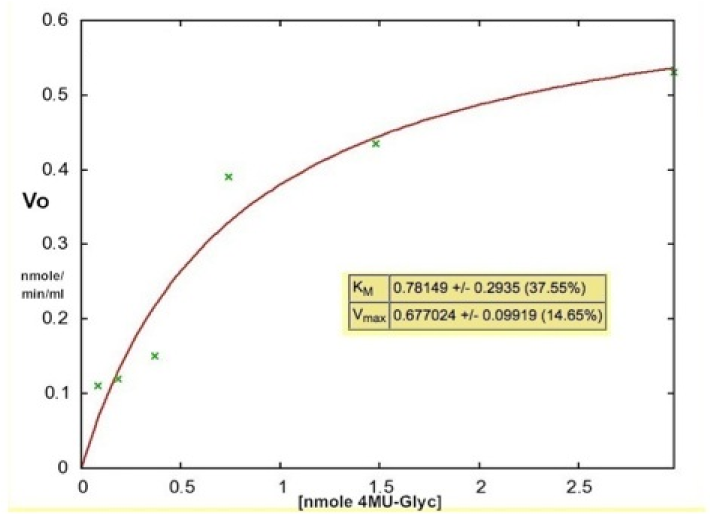
4MU-Glc Km & Vmax for barley >50kD are bar graphs showing the Km, Vmax and IC50 for the 50-100kD fraction for 4-MU-Glc, glycogen, maltose and 4-MU-Glc/glycogen.

**Figure 8.**
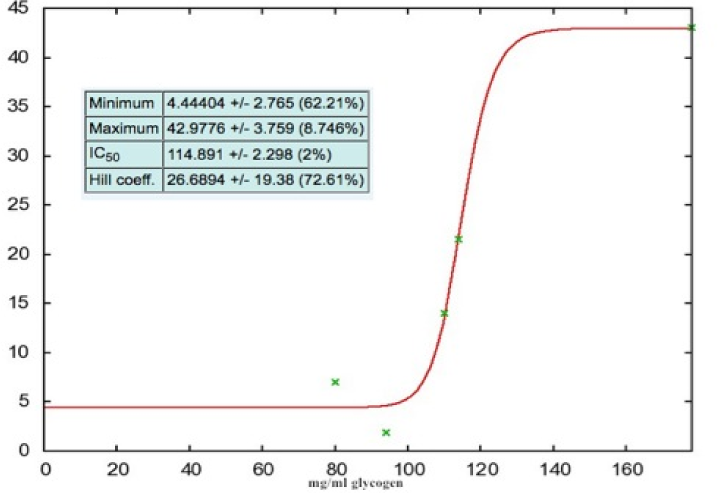
IC50 for glycogen v.4MU-Glc with Robust bGAA are bar graphs showing the Km, Vmax and IC50 for the 50-100kD fraction for 4-MU-Glc, glycogen, maltose and 4-MU-Glc/glycogen.

**Figure 9.**
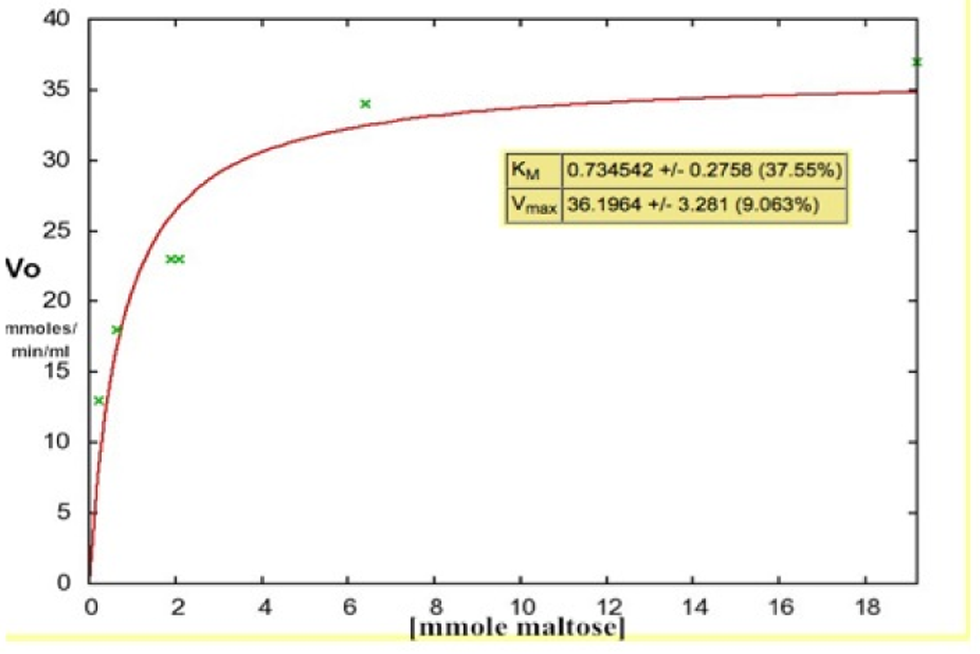
Km & Vmax OF maltose for Barley >50kD are bar graphs showing the Km, Vmax and IC50 for the 50-100kD fraction for 4-MU-Glc, glycogen, maltose and 4-MU-Glc/glycogen.

**Figure 10.**
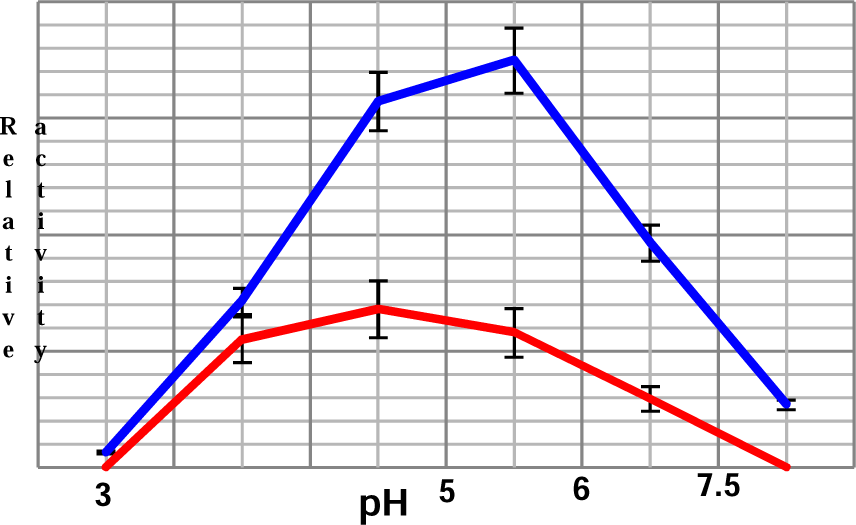
pH Optima for Robust Barley 50-100kD for 4-MU-Glc or Glycogen is a bar graph showing the pH optima for 4-MU-Glc (pH 5.5) and glycogen (pH 4.5) for the 50-100kD fraction.

#### A9-Affect of stomach and small intestinal environment (79,80)

To mimic the stomach and small intestine (SI) environment, we exposed the **L-GGB** to physiologic conditions/times to pepsin (stomach) and trypsin/chymotrypsin (small intestine). GAA is stable at low pH. We exposed **L-GGB** to 300 μg/ml pepsin at pH 4.0 and then trypsin at 800 μg/ml-pH 6.5/chymotrypsin at 700 μg/ml-pH 6.5 for 60 minutes each at 37°C. None of the conditions had any affect on bGAA activity from **L-GGB** thus demonstrating that conditions in the digestive tract do not affect **L-GGB**.

#### A10-Nutritional facts

We had nutritional ingredients determined at Exact Scientific Services for a 100 ml serving (**Table 2**)(Inc.www.exactscientific.com). The pH was 4.1.

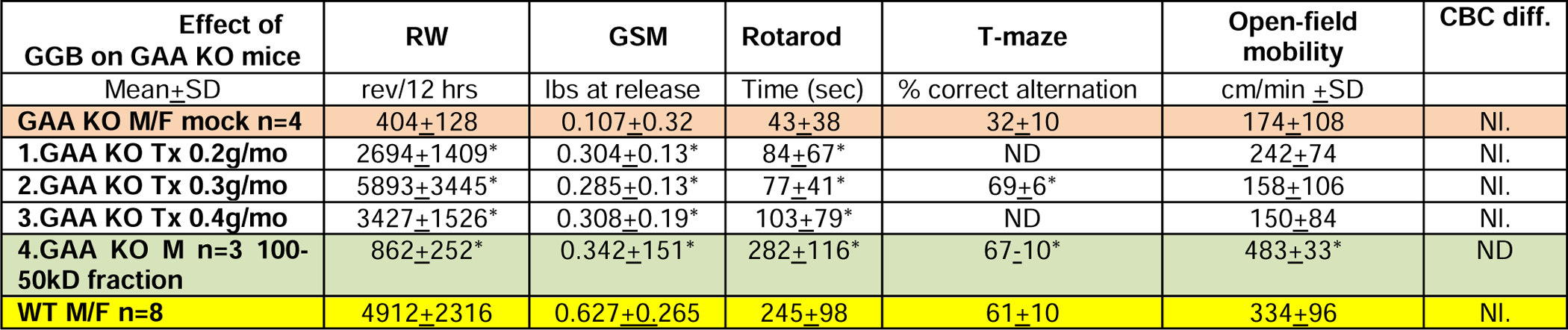
Table 2.

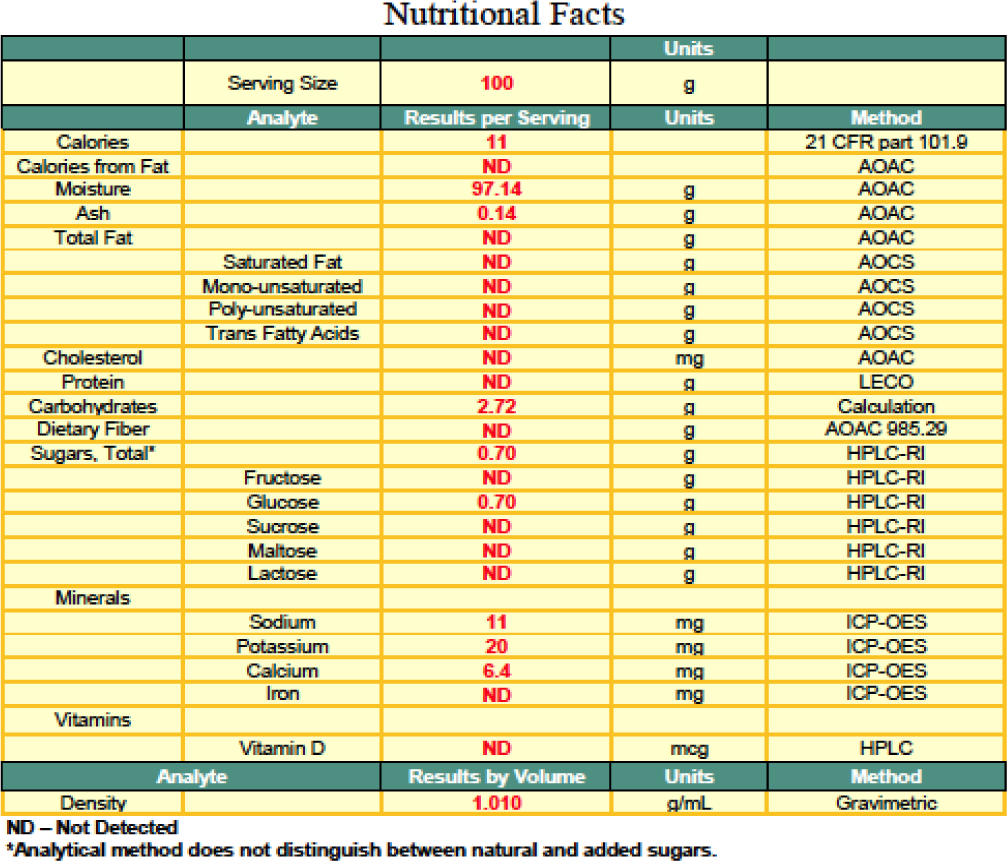
Table 3.

#### A11-Ammonium sulfate precipitation

bGAA was precipitated at 60-80% saturation that will be useful for purification and was similar to hGAA (**82,83**).

### B-*in vitro* studies

#### B1-Uptake of L-GGB by white blood cells from human and GAA KO mice

Uptake of **L-GGB** by white blood cells from human and GAA KO mice were tested (±5mM mannose-6-phosphate, blocks uptake by M6P receptor, **98-100**) and as a positive control-placental hGAA. Human WBCs exposed to **L-GGB** or placental hGAA had higher ratios of GAA/NAG compared to mock treated cells (**Figure 11**). bGAA was also taken up by mouse GAA KO WBCs and uptake was blocked by mannose-6-phosphate similar to placental hGAA (mean±SD) (**Figure 12**). These experiments indicated that human and mouse WBCs can take up bGAA.

**Figure 11.**
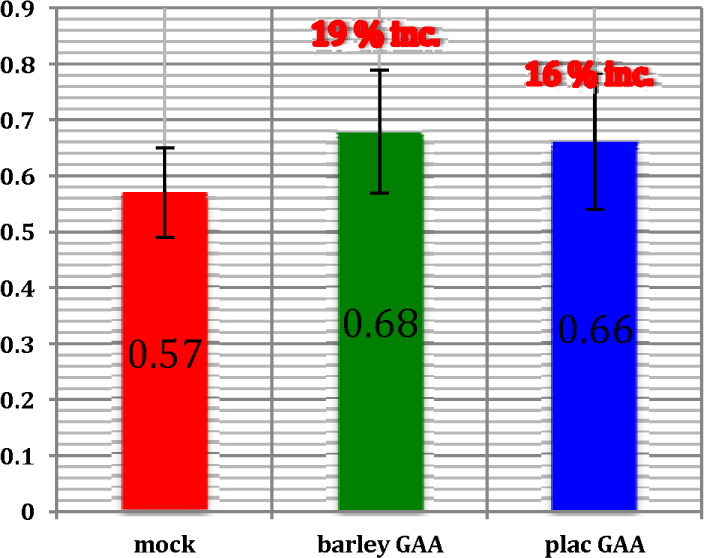
Human WBCs GAA/NAG exposed to day 7 germinated barley GAA or placental GAA is a bar graph showing the GAA/NAG ratio after uptake of L-GGB by isolated human white blood cells (WBCs) compared to uptake of placental human GAA as a positive control (mean+SD).

**Figure 12.**
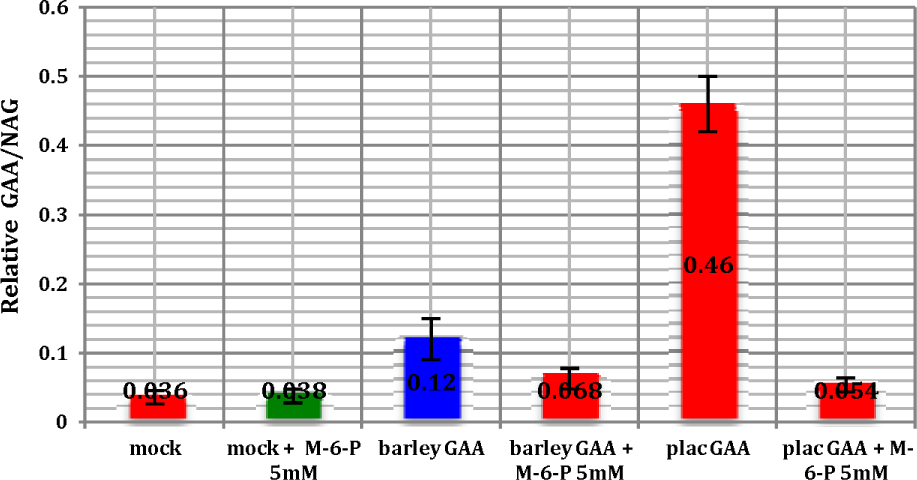
GAA KO mouse WBCs GAA/NAG exposed to day 7 germinated barley GAA or placental GAA + 5mM mannose-6-phosphate is a bar graph showing GAA KO mice white blood cells and their GAA/NAG ratio after uptake of L-GGB or placental human GAA treated with and without 5mM M6P (which blocks uptake by M6P receptor). bGAA was taken up by mouse GAA KO WBCs and blocked by M6P similar to human placental GAA (mean+SD).

#### B2-Uptake of L-GGB or 50-100kD fraction in a human PD fibroblast and lymphoid cell lines

Human lymphoid (GM6314, GM13793, GM14450) or fibroblast (GM4912, GM1935, GM3329) cell lines from infantile or adult PD exposed to varying amounts of **L-GGB** or 50-100kD fraction or placental hGAA or rhGAA (R&D Systems) for 48 hours and assayed. At 48 hours, equivalent concentrations showed similar uptake and increases in GAA (mean±SEM)(**Figures 13 and 14**).

**Figure 13.**
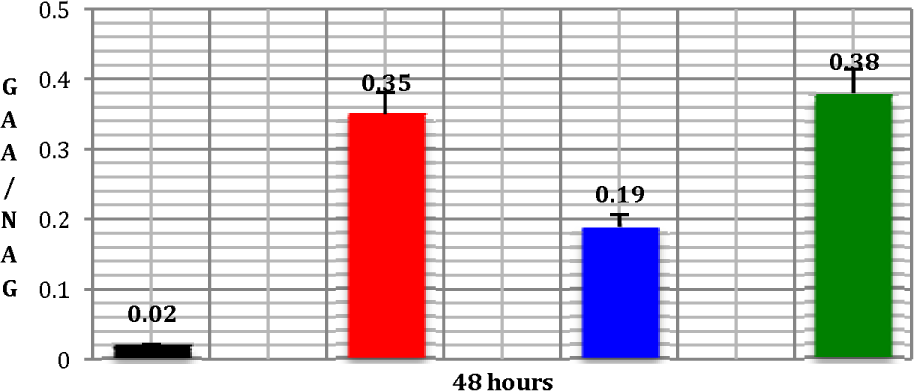
Human PD Lymphoid Lines Exposed to Mock Treated or bGAA 100-50kD or plac GAA or rhGAA for 48 Hours is a bar graph showing human PD lymphoid lines exposed to mock treatment, bGAA 50-100kD fraction, human placental GAA or rhGAA for 48 hours and assayed for GAA. All treatments were significant at p = <0.05.

**Figure 14.**
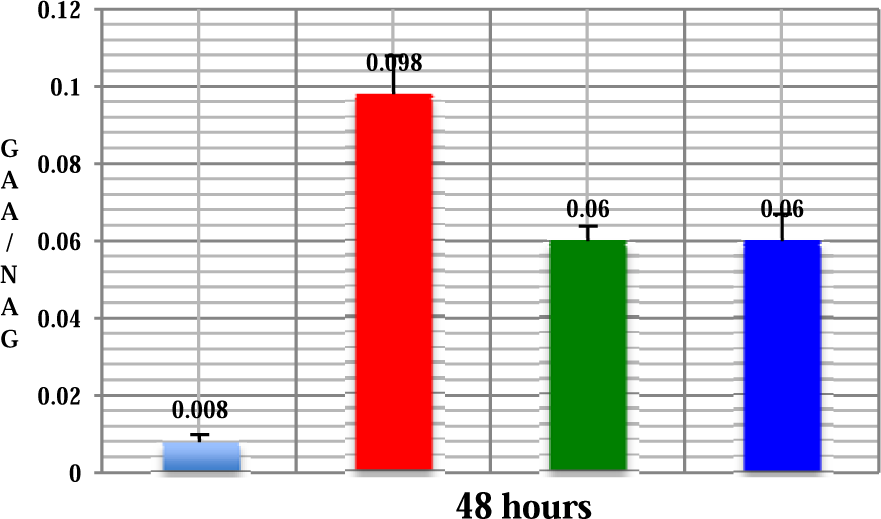
Human PD fibroblasts exposed to either mock or L-GGB 50-100kD or plac GAA or rhGAA for 48 hours is a bar graph showing human fibroblast cell lines human fibroblast (GM4912, GM1935, GM3329) cell lines from infantile or adult PD exposed to mock treatment, **L-GGB** 50-100kD fraction, human placental GAA or rhGAA for 48 hours and assayed for GAA. All treatments were significant at p = ≤0.05.

#### B.3-Uptake of L-GGB or 50-100kD fraction in a human PD myoblast cell line

A human Pompe disease myoblast cell line (homozygous for the IVS1 c.-32–13t>g)(**102**) was exposed to various amounts of **L-GGB** or 50-100kD fraction for 48 hours and assayed. Mock treated GAA and normal skeletal muscle cells were controls plus cells treated with rhGAA. **Figure 15** shows both **L-GGB** and the 50-100kD bGAA fraction increased GAA to 19-51% of normal (mean±SD)(**Figure 15**). All treatments were significant at p = ≤0.05. Simultaneously, we measured cell proliferation with the Cayman MTT assay and found that **L-GGB** slightly reduced cell growth by <10%.

**Figure 15.**
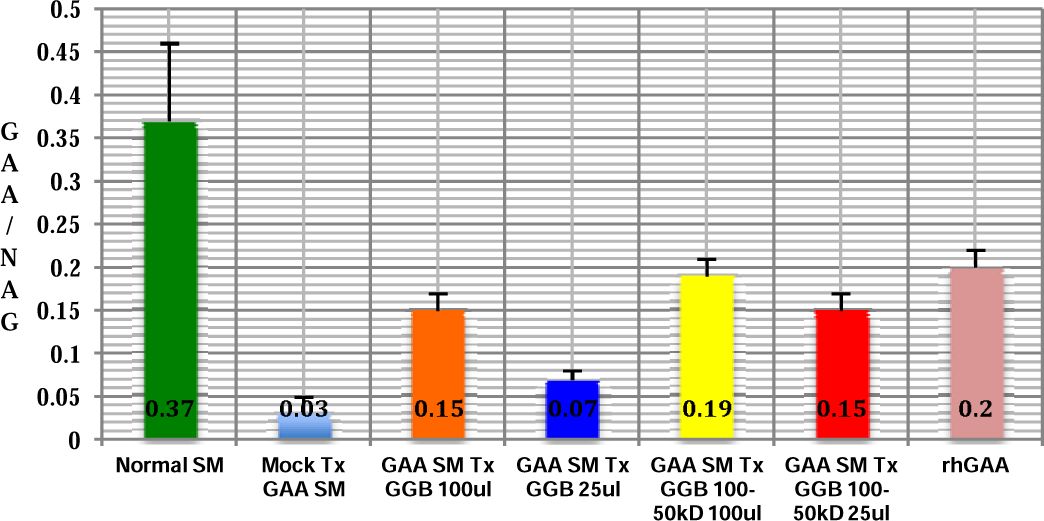
Human normal and PD myoblast cells exposed to L-GGB or L-GGB 50-100kD or rhGAA is a bar graph showing human normal and PD myoblast cells exposed to various amounts of L-GGB or L-GGB 50-100kD or rhGAA for 48 hours and assayed for GAA. We found an increased GAA to almost 50% of normal (mean+SD). All treatments were significant at p = <0.05.

### C-*in vivo* studies

#### C1-A time and dose-response in GAA KO mice

GAA KO mice (∼6 months old) were fed daily with germinated (7-day germination) barley seeds in amounts of either 0, 1, 3, 6, 12, 24 or 48 seeds for two weeks at each level. Upon consuming bGAA, GAA KO mice showed a reversal in hind-limb muscle weakness as assessed by increased running wheel (RW) activity that was dose dependent (**Figure 16**). GAA KO mice treated daily with un-germinated seeds showed no increase in RW activity.

**Figure 16.**
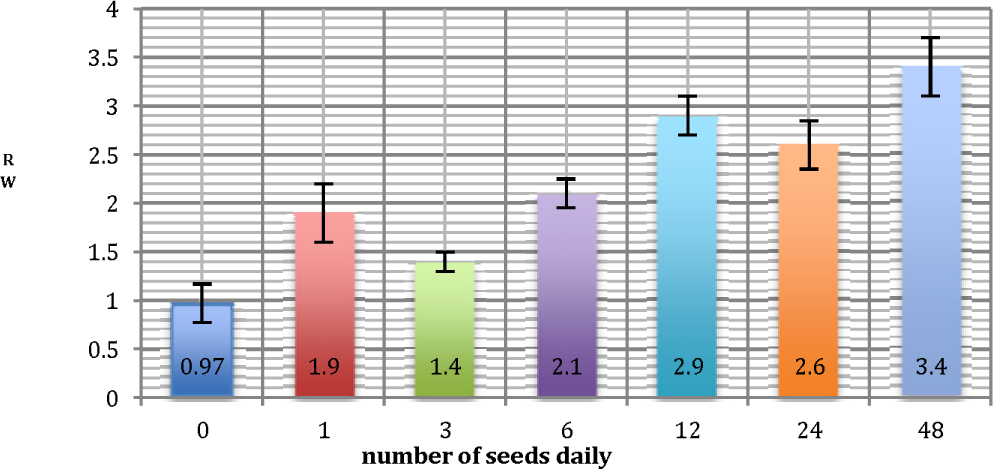
Ratio of RW activity in treated and mock GAA KO mice feed different amounts of germinated barley seeds daily for 2 weeks each level is a bar graph showing the ratio of RW activity in treated and mock GAA KO mice feed different amounts of germinated barley seeds of either 0, 1, 3, 6, 12, 24 or 48 seeds for two weeks at each level.

#### C2-Assessment of reversal of clinical phenotypes

GAA KO mice treated daily with **GGB** on peanut butter/mouse chow in a petri dish at 0.2g/mouse for 3 weeks (#1) followed by feeding **GGB** at 0.3g/mouse for 3 weeks (#2) and then **GGB** at 0.4g/mouse for 3 weeks (#3). After 3 weeks of feeding at each dose, mice were evaluated for motor activity by running wheel (RW); fore-limb muscle strength by grip-strength meter (GSM), motor coordination/balance with a rotarod, open-field mobility by 5-min video, CBC (complete blood count) differentials and spontaneous learning with a T-maze (**Table 2**). The * indicates p=<0.05 when compared to pre-treatment data (1-tail, 2-equal variance). For all parameters and doses, improvements from pre-treated GAA KO was significant except for open field mobility and were approaching WT levels. We next treated mice daily with 0.3 ml of the 50-100kD from **L-GGB** (in mouse food/peanut butter) for 3 weeks and assessed clinical phenotypes. Thus, the 50-100kD fraction showed significant improvements in the phenotypes demonstrating that bGAA may be the major factor for these improvements.

The T-maze test to evaluate spontaneous learning showed an increase in the % correct alternation for **GGB** fed GAA KO mice compared to control GAA KO mice. Thus, statistically significant improvements from pre-treated GAA KO mice for RW, GSM, Rotarod and T-maze were observed for all three doses of **GGB** administration. CBC differentials of treated GAA KO animals were identical to pre-treatment and treated GAA KO plus WT mice. Spontaneous alternation is used to assess the cognitive ability of rodents to choose one of the 2 goal arms of the T-maze The advantage of a free choice procedure is that hippocampal or lesioned animals often develop a side preference and scores below 50%. Controls generally achieve at least 60-80% correct alternation (**106**). Spontaneous alternative learning for cognitive ability was assessed in the T-maze in both male and female GAA KO mice and WT mice from 2-9 mo of age. It was found that deficiency in spontaneous learning appeared by 2-3 months in male and 3-4 months in female GAA KO mice. All conditions were significant (p<0.05).

These experiments revealed that bGAA from barley in the form of **GGB** significantly improved motor and cognitive neurological functions as assessed by multiple different tests in GAA KO mice in a dose dependent manner. Running wheel times, grip strength meter, Rotarod performance tests, T-maze and spontaneous alternation tests were all significantly improved compared to GAA KO mice that did not receive **GGB**.

#### C3-Assessment of cognitive ability

by spontaneous alternation for rodents to choose one of 2 arms of a T-maze (**106**). The advantage of a free choice procedure is that hippocampal or lesioned animals often develop a side preference and scores below 50%. Controls achieve 60-80% correct alternation. We assessed spontaneous alternative learning for cognitive ability in the T-maze in male and female GAA KO mice and WT mice from 2-9 months of age. We found that deficiency in spontaneous learning appeared by 3 months in males and 4 months in female GAA KO mice. All conditions were significant (p≤0.05)(**Figure 17**). We found that deficiency in spontaneous learning appeared by 2-3 months in male and 3-4 months in female GAA KO mice.

**Figure 17.**
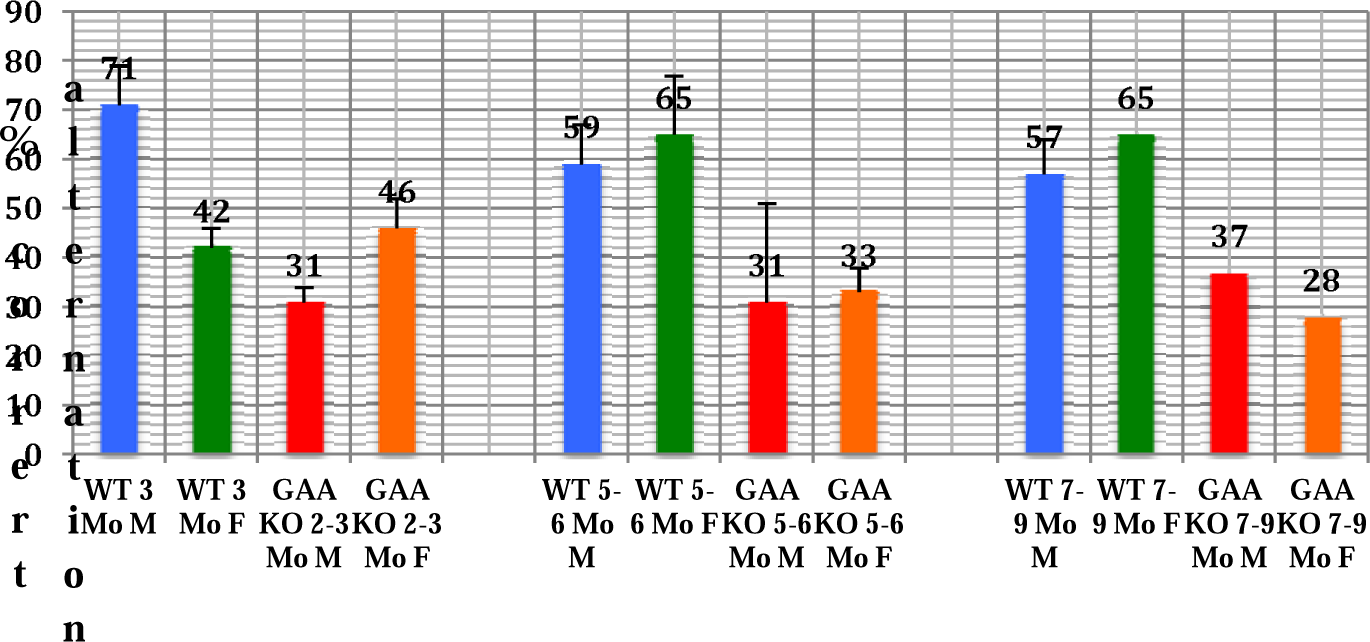
T-maze over time for WT and GAA KO males and females is a bar graph of spontaneous alternative learning for cognitive ability in the T-maze in male and female GAA KO mice and WT-129/C57 mice from 2-9 months of age. Deficiency in spontaneous learning appeared by 3 months in males and 4 months in female GAA KO mice. All conditions were significant (p≤0.05).

#### C4-IL6 baseline for WT and GAA KO mice

in serum with the PeproTech’s pro-inflammatory cytokine IL6 ELISA was measured every 2 weeks for 3 months, the IL6 levels for all were below detection of the lowest standard of 16pg/ml.

#### C5-Steady-state GAA/NAG levels in tissues

We measured steady-state GAA/NAG from the mice in **Table 4**. The % of WT activity in tissues ranged from 4 to 53%.

Table 4.

| **Table 4** | liver | heart | Hind-limb tibialis anterior muscle | kidney | lung | diaphragm | brain | SI-D | SI-J | SI-I | spleen | pancreas |
| --- | --- | --- | --- | --- | --- | --- | --- | --- | --- | --- | --- | --- |
| GAA KO mock | 0.073 | 0.06 | 0.018 | 0.085 | 0.094 | 0.045 | 0.012 | 0.4 | 0.94 | 0.71 | 0.015 | 0.084 |
| GAA KO Tx 0.3ml >50kD L-GGB | 0.44 | 0.35 | 0.08 | 0.2 | 0.164 | 0.09 | 0.026 | 0.96 | 0.84 | 0.76 | 0.104 | 0.019 |
| WT-129/C57 | 4.3 | 3.1 | 0.71 | 1.67 | 0.31 | 0.37 | 0.73 | 2.68 | 3.47 | 3.3 | 1.06 | 0.67 |
| % WT | 10 | 11 | 11 | 11 | 53 | 12 | 4 | 37 | 21 | 22 | 10 | 3 |

## DISCUSSION

Currently, there is no effective treatment or cure for people living with PD. Sanofi-Genzyme, Inc. using a rhGAA (alglucosidase alfa) secreted from a CHO cell line has demonstrated moderate success in patients (**27**), however yearly costs are very high. Although alglucosidase alfa has been a wonderful first step in treating PD it has revealed subtle aspects that must be dealt with for successful treatment (**18,47–54**). ERT usually begins when the patients are symptomatic, however, all the secondary problems are already present which are further compounded by issues with alglucosidase alfa uptake in muscle (**Table 5**)(**18**). Lysosomal enzymes (such as GAA) are targeted to the lysosome by a M6P recognition sequence that is exposed by posttranslational modification in the Golgi that may be the mechanism that extracellular GAA can be recycled and targeted back to the lysosomes. This mechanism will potentially allow rhGAA to be delivered to the cells or tissues and directed to the lysosome. However, some GAA may be taken up or recycled by endocytosis or a mannose-6-phosphate independent mechanism (**22–25**). In skeletal muscle there are a low abundance of the cation-independent M6P receptors (CI-MPR) compounded by low blood flow plus low affinity of the CHO-produced alglucosidase alfa for the CI-MPR. Alglucosidase alfa is delivered to the lysosome by receptor-mediated endocytosis after binding of the M6P-glycan to the CI-MPR. Alglucosidase alfa has one M6P/enzyme (**107–109**) and 1,000-fold lower affinity (**110**). With alglucosidase alfa, the high level of infused enzyme is transient, leaving the patients with no therapeutic enzyme for the remaining 13 days. Thus, oral formulations of ground tobrhGAA seeds will allow patients to ingest daily in single or multiple doses conveniently at home to sustain a therapeutic dose of enzyme activity and to improve the quality of life.

**Table 5.**
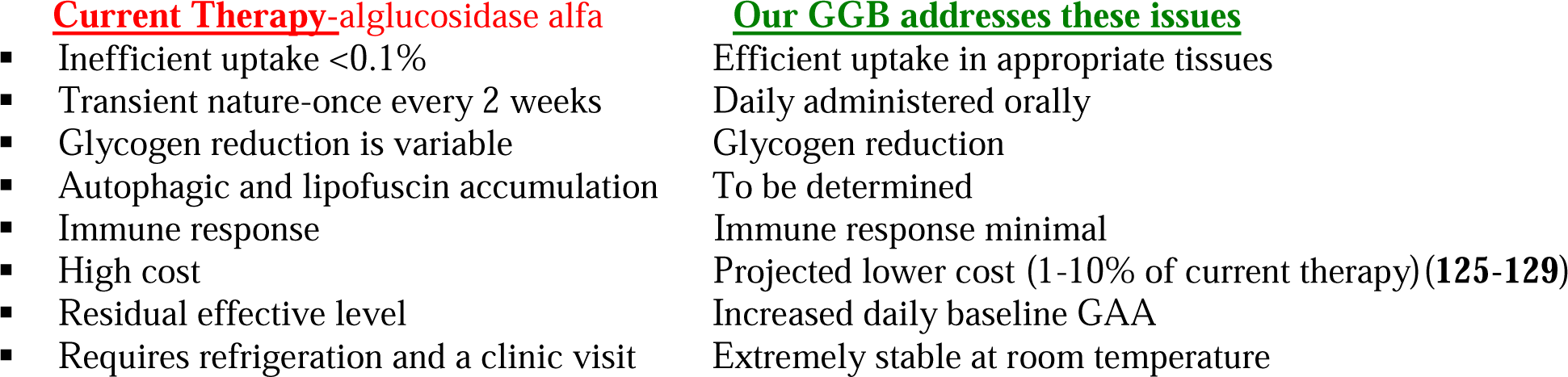
Comparison of alglucosidase alfa and GGB.

Investigators are studying the role of autophagy (**18,111–113**), an intracellular system for delivering portions of cytoplasm and damaged organelles to lysosomes for degradation/recycling in PD and the reduction of glycogen in skeletal muscle. A GAA and glycogen synthase 1 (*GYS1*) KO mouse exhibited a profound reduction of glycogen in the heart and skeletal muscles, a decrease in lysosomal autophagic build-up and correction of cardiomegaly. The abnormalities in glucose metabolism were corrected in the double-KO mice that demonstrate long-term elimination of muscle glycogen synthesis leads to a significant improvement of structural, metabolic and functional defects and a new perspective for treatment of PD (**17**). Lim et al. (**113**) reactivated mTOR in PD mice by TSC knockdown resulted in the reversal of muscle atrophy, removal of autophagic buildup plus aberrant mTOR signaling can be reversed by arginine alone (**112–114**). Jung et al. (**115,116**) produced and characterized a rhGAA in a transgenic rice cell suspension culture with high-mannose glycans and was similar to the CHO derived hGAA.

Plant cell cultures have emerged as a promising platform for the production of biopharmaceutics due to their cost-effectiveness, safety, ability to control the cultivation and secrete products into culture medium. However, the use of this platform is hindered by the generation of plant-specific *N*-glycans, the inability to produce essential *N*-glycans for cellular delivery of biopharmaceutics and low productivity. Sariyatun et al. (**117**) producted a hGAA for ERT with a paucimannose structure-Man3GlcNAc2 (M3), without the plant-specific *N*-glycans by glycoengineered in *Arabidopsis alg3* cell culture.

Seeds may be a better vehicle for Oral-ERT for lysosomal diseases, such as PD, compared to intravenous systemic delivery. Seeds and not other plant tissues and organelles, contain the metabolic machinery necessary for correct glycosylation, processing, phosphorylation and synthesis of complex enzymes and proteins, enzymes or proteins plus are protected or shielded from digestion in the stomach and small intestine and can be administered daily in single and multiple doses and seeds provide long-term stable storage of recombinant enzymes. A number of biotechnology companies have tried to mass produce a rhGAA from different platforms. A consensus at a 2019 US Acid Maltase Deficiency conference at San Antonio, TX, suggested that a multi-pronged approach including gene therapy, diet, exercise, etc. must be evaluated for a successful treatment of Pompe disease. Prompted by studies of oral formulations of edible tissues in broccoli sprouts, corn/maize, pea, rice, tobacco and tomato for pharmaceutical applications including vaccination to induce effective mucosal immune tolerance and immune reactions, GI, bacterial/viral infections, allergies, asthma, diabetes, endocrine-associated diseases, hypersensitivity, elevated BP, cholera, leucopenia, cancer and RA (**118–138**), we hypothesized that rhGAA from edible plant tissues offers an cost-effective, innovative and safe strategy for Oral-ERT for PD. Previously, we investigated the potential of genetically engineered “edible plant tissues” as an alternative large-scale production system that overcomes the high cost of producing rhGAA in CHO cells. Oral-ERT can be safer than infusion and be ingested in a pill/capsule format at frequent intervals daily to maintain a therapeutic dose of enzyme to achieve long-term clinical efficacy (**139–143**). Thus, to provide a less expensive alternative, we generated a rhGAA produced in tobacco seeds for ERT of PD. We found that the tobrhGAA compared favorably and was superior on many aspects to alglucosidase alfa.

Using plant cells to make biopharmaceuticals is not a new idea. The biopharmaceutical industry has been slow, but not entirely resistant, to embrace plant-based manufacturing, which is also known as molecular pharming. Current biopharmaceuticals are prohibitively expensive due to production in expensive fermenters, purification, cold storage, short shelf-life and sterile delivery methods ($140 billion in 2013, exceeds the GDP of >75% of countries) making them unaffordable. There are many different ways in which plants or plant cells could be used in biopharmaceutical manufacturing, ranging from using outdoor, field-grown plants to growing plant cells in liquid suspension in bioreactors, but also including intermediate approaches such as plants grown in greenhouses or in indoor vertical agriculture facilities. Urgent unmet medical needs of metabolic or genetic disorders to answer high levels of proteins in edible cells, storage at ambient temperature for years, scale-up of production in cGMP facilities or as dietary supplements will make them affordable (**144**). Thus, we wanted to identify alternate forms of GAA from various sources to supplement or provide a safe, cheap and tolerable ERT for PD. Literature searches lead us to germinated barley as a possible source. We evaluated a detailed study in GAA KO mice of utility of **GGB** or **L-GGB** as ERT or as a supplement to Myozyme. Our innovation is two-fold: (i) **GGB** as a dietary supplement for Oral-ERT and (ii) oral delivery of an active bGAA in **GGB** or **L-GGB** for the long-term treatment of PD. In contrast to the current ERT of an infusion every 2 weeks in a clinical setting, a dietary supplement with **GGB** or **L-GGB** allows patients to ingest it daily in single or multiple doses conveniently at home. Successful demonstration in this grant will have significant clinical applications and shift current clinical practice paradigm by offering a safe and affordable lifelong treatment. Although Myozyme has been a wonderful first step in treating PD, it revealed subtle aspects that must be dealt with for successful treatment (**Table 5**). These experiments revealed that GAA from barley in the form of **GGB** or **L-GGB** significantly improved motor and cognitive neurological functions as assessed by multiple different tests in GAA KO mice in a dose dependent manner. Running wheel times, grip strength meter, Rotarod performance tests, T-maze and spontaneous alternation tests were all significantly improved compared to GAA KO mice that did not receive **GGB** or **L-GGB**. These results were unexpected given that the bGAA has only approximately 39% sequence identity and 55% sequence similarity over a limited portion of the protein compared to the human GAA protein but has complete homology at the active site and surrounding amino acids. The bGAA was shown to be taken up by both human and murine white blood cells in a similar fashion as human rhGAA. Furthermore, the oral delivery of bGAA was shown to be efficacious in improving multiple indices of skeletal muscle and cognitive neurological function that were affected by the lysosomal storage disorder caused by *GAA* deficiency. **GGB** or **L-GGB** has many advantages over the current treatments for PD. The cost of producing **GGB** or **L-GGB** is very low compared to the costs of producing rhGAA and is extremely stable at room temperature. **GGB** or **L-GGB** is easily administered and does not require a visit to a physician’s office or clinic or intravenous infusion as required by the current treatment. Furthermore, **GGB** or **L-GGB** showed a reduced or absent immunological reaction and an ability to reverse neurological problems.

Estimates of uptake of alglucosidase alfa (low M6P content) are ∼0.1% or 1.4mg/70kg LOPD for the infused 1.4g dose. **L-GGB** contains ∼0.04mg/ml or ∼30-50 ml can taken daily to be equivalent to the 1.4mg/70kg LOPD if uptake is 100% efficient. Based upon *in vitro* data for uptake by PD fibroblasts (**36**), we estimate the % accumulation in daily treatment with a 20% decline daily that by day 6 there was 3.2 times the daily dose (**Figure 18**).

**Figure 18.**
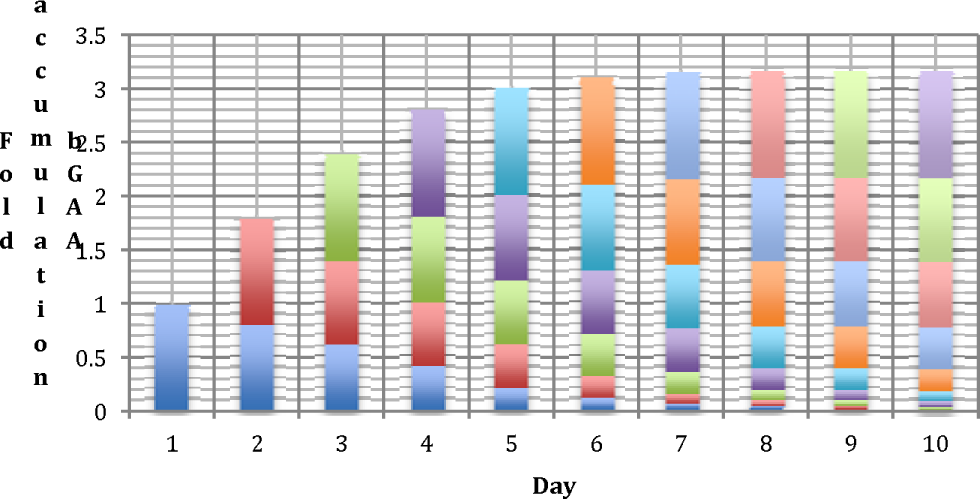
Predicted Accumulation of bGAA Taken upDaily Orally is a bar graph showing predicted accumulation of bGAA taken daily orally of L-GGB which contains ∼0.04mg/ml or ∼30-50 ml can taken daily to be equivalent to the 1.4mg/70kg LOPD alglucosidase alfa (low M6P content) of ∼0.1%. Based upon in vitro data for uptake by PD fibroblasts, we estimate the % accumulation in daily treatment with a 20% decline daily that by day 6 there was 3.2 times the daily dose.

Successful demonstration of Oral-ERT with **GGB** or **L-GGB** will have significant clinical applications and shift current clinical practice paradigm by offering a safe, stable at room temperature and affordable lifelong Oral-ERT for PD in the USA and world-wide especially in areas where access to a clinical setting for biweekly IV administration of Myozyme is lacking. We estimate that an adults living with PD patient will need to take 3-4 doses daily at a cost of <1% of Myozyme depending on how it is branded and marketed to maintain a sustained GAA level of 5-10% of normal.

## Abbreviations

AMD: acid maltase deficiency
rhGAA: recombinant GAA
GAA: acid maltase
ERT: enzyme replacement therapy
Oral-ERT: oral enzyme replacement therapy
Pompe disease: PD
GGB: ground germinated barley
L-GGB: liquid ground germinated barley
bGAA: barley GAA
GA: gibberellic acid
M6P: mannose-6-phosphate.

## ACKNOWLEDGEMENTS

No artificial intelligence software or programs were used in preparation of the manuscript.

Data is protected by the patent-“Methods and compositions for treatment of lysosomal storage”. Inventors: Frank Martiniuk, Elena Arvanitopoulos, Angelo Kambitsis, Kam-Meng Tchou-Wong and John Arvanitopoulos, Assignee-JME Group, Inc. non-provisional patent (17/404,680), Feb, 2022.

## REFERENCES

1. Slonim A, Bulone L, Ritz S, Goldberg T, Chen A, Martiniuk F. Identification of two subtypes of infantile acid maltase deficiency: Evaluation of twenty-two patients and review of the literature. J Pediatrics. 2000;137:283–5.

2. Engel AG. Acid maltase deficiency. In Engel AG, Franzini-Armstrong C, eds. Myology, N.Y., McGraw Hill, Inc. p. 1533–1553, 1994.

3. Reuser AJJ, Kroos M, Willemsen R, Swallow D, Tager JM, Galjaard H. Clinical diversity in glycogenosis type II. J Clin Invest. 1987;79:1689–1699.

4. Beratis N, LaBadie GU, Hirschhorn R. Genetic heterogeneity in acid alpha glucosidase deficiency. Am J Hum Genet. 1983;35:21–33.

5. Tagers JM, Oude Elferink RPJ, Reuser AJJ, Kroos M, Ginsel LA, Franzen JAM, Klumperman J. Alpha glucosidase deficiency; Pompe’s disease. Enzyme. 1987;38:280–285.

6. van den Hout HM, Hop W, van Diggelen OP, Smeitink JA, Smit GP, Poll-The BT, Bakker HD, Loonen MC, de Klerk JB, Reuser AJ, van der Ploeg AT. The natural course of infantile Pompe’s disease: 20 original cases compared with 133 cases from the literature. Pediatrics. 2003;112:332–340.

7. Winkel LP, Hagemans ML, van Doorn PA, Loonen MC, Hop WJ, Reuser AJ, van der Ploeg AT. The natural course of non-classic Pompe’s disease; a review of 225 published cases. J Neurol. 2005;252:875–884.

8. Engel AG, Gomez MR, Seybold ME, Lambert EH. The spectrum and diagnosis of acid maltase deficiency. Neurology. 1973;23:95–106.

9. Mehler M, DiMauro S. Residual GAA activity in late-onset acid maltase deficiency. Neurol. 1997;27:178–184.

10. La Badie GU. Biochemical and immunologic studies of acid alpha glucosidase deficiency, a genetically heterogeneous, inherited neuromuscular disease. (Ph.D. Thesis, City University of New York, Mt. Sinai Hospital, 1986).

11. Raben N, Ralston E, Chien YH, Baum R, Schreiner C, Hwu WL, Zaal KJ, Plotz PH. Differences in the predominance of lysosomal and autophagic pathologies between infants and adults with Pompe disease: implications for therapy. Mol Genet Metab. 2010;101:324–331.

12. Martiniuk F, Chen A, Mack A, Codd W, Hanna B, Arvanitopoulos E, Slonin A. Carrier frequency for glycogen storage disease type II in New York and estimates of affected individuals born with the disease. Am J Med Genet. 1998;79:69–72.

13. Park K.S. Carrier frequency and predicted genetic prevalence of Pompe disease based on a general population database. Molecular Genetics and Metabolism Reports. 2021;27:100734–42.

14. Reuser AJJ, Kroos M, Elferink RPJO, Tager JM. Defects in synthesis, phosphorylation, and maturation of acid alpha glucosidase. J Biol Chem. 1985;260:8336–8341.

15. Strothotte S, Strigl-Pill N, Grunert B, Kornblum C, Eger K, Wessig C, Deschauer M, Breunig F, Glocker FX, Vielhaber S, Brejova A, Hilz M, Reiners K, Müller-Felber W, Mengel E, Spranger M, Schoser B. Enzyme replacement therapy with alglucosidase alfa in 44 patients with late-onset glycogen storage disease type 2:12-month results of an observational clinical trial. J Neurol. 2010;257:91–97.

16. van der Ploeg AT, Clemens PR, Corzo D, Escolar DM, Florence J, Groeneveld GJ, Herson S, Kishnani PS, Laforet P, Lake SL, Lange DJ, Leshner RT, Mayhew JE, Morgan C, Nozaki K, Park DJ, Pestronk A, Rosenbloom B, Skrinar A, van Capelle CI, van der Beek NA, Wasserstein M, Zivkovic SA. A randomized study of alglucosidase alfa in late-onset Pompe’s disease. N Engl J Med. 2010;362:1396–1406.

17. Douillard-Guilloux G, Raben N, Takikita S, Ferry A, Vignaud A, Guillet-Deniau I, Favier M, Thurberg BL, Roach PJ, Caillaud C, Richard E. Restoration of muscle functionality by genetic suppression of glycogen synthesis in a murine model of Pompe disease. Human Molecular Genetics. 2010;19:684–696.

18. Lim J, Li L, Raben N. Pompe disease: from pathophysiology to therapy and back again. Frontiers in Aging Neuroscience. 2014;6:1–14.

19. van Gelder CM, Hoogeveen-Westerveld M, Kroos MA, Plug I, van der Ploeg AT, Reuser AJJ. Enzyme therapy and immune response in relation to CRIM status: the Dutch experience in classic infantile Pompe disease. J Inherit Metab Dis. 2015;38:305–314.

20. Lacaná E, Yao LP, Pariser AR, Rosenberg AS. The role of immune tolerance induction in restoration of the efficacy of ERT in Pompe disease. Am J Med Genet Part C Semin Med Genet. 2012;160C:30–39.

21. Bali DS, Goldstein JL, Banugaria S, Dai J, Mackey J, Rehder C, Kishnani PS. Predicting cross-reactive immunological material (CRIM) status in Pompe disease using GAA mutations: Lessons learned from 10 years of clinical laboratory testing experience. Am J Med Genet Part C Semin. 2012;160C:40–49.

22. Joseph A, Munroe K, Housman M, Garman R, Richards S. Immune tolerance induction to enzyme-replacement therapy by co-administration of short-term, low-dose methotrexate in a murine Pompe disease model. Clinical and Experimental Immunology. 2013;152:138–146.

23. Banugaria SG, Prater SN, McGann JK, Feldman JD, Tannenbaum JA, Bailey C, Gera R, Conway RL, Viskochil D, Kobori JA, Rosenberg AS, Kishnani PS. Bortezomib in the rapid reduction of high sustained antibody titers in disorders treated with therapeutic protein: lessons learned from Pompe disease. Genet Med. 2013;15:123–131.

24. Berrier KL, Kazi ZB, Prater SN, Bali DS, Goldstein J, Stefanescu MC, Rehder CW, Botha EG, Ellaway C, Bhattacharya K, Tylki-Szymanska A, Karabul N, Rosenburg AS, Kishnani PS. CRIM-negative infantile Pompe disease: characterization of immune responses in patients treated with ERT monotherapy. Genet Med. 2015;17:912–918.

25. Banugaria SG, Prater SN, Ng YK, Kobori JA, Finkel RS, Ladda RL, Chen YT, Rosenberg AS, Kishnani PS. The impact of antibodies on clinical outcomes in diseases treated with therapeutic protein: Lessons learned from infantile Pompe disease. Genetics in Medicine. 2011;13:729–736.

26. Messinger YH, Mendelsohn NJ, Rhead W, Dimmock D, Hershkovitz E, Champion M, Jones SA, Olson R, White A, Wells C, Bali D, Case LE, Young SP, Rosenberg AS, Kishnani PS. Successful immune tolerance induction to enzyme replacement therapy in CRIM–negative infantile Pompe disease. Genet Med. 2012:14:135–142.

27. Al Khallaf HH, Geffrard J, Botha E, Ali Pervaiz M. CRIM-negative Pompe disease patients with satisfactory clinical outcomes on enzyme replacement therapy. JIMD Reports. 2012;192:133–7.

28. Brooks DS. Immune response to enzyme replacement therapy in lysosomal storage disorder patients and animal models. Molecular Genetics and Metabolism. 1999;68:268–275.

29. Kishnani PS, Goldenberg PC, DeArmey SL, Heller J, Benjamin D, Young S, Bali D, Smith SA, Li JS, Mandel H, Koeberl D, Rosenberg A, Chen YT. Cross-reactive immunologic material status affects treatment outcomes in Pompe disease infants. Molecular Genetics and Metabolism. 2010;99:26–33.

30. de Vries JM, van der Beek N, Kroos MA, Özkan L, van Doorn PA, Richards SM, Sung CCC, Brugma JDC, Zandbergen AAM, van der Ploeg AT, Reuser AJJ. High antibody titer in an adult with Pompe disease affects treatment with alglucosidase alfa. Molecular Genetics and Metabolism. 2010;101:338–345.

31. Patel TT, Banugaria SG, Case LE, Wenninger S, Schoser B, Kishnani PS. The impact of antibodies in late-onset Pompe disease: A case series and literature review. Molecular Genetics and Metabolism. 2012;106:301–309.

32. Wang J, Lozier J, Johnson G, Kirshner S, Verthelyi D, Pariser A, Shores E, Rosenberg A. Neutralizing antibodies to therapeutic enzymes: considerations for testing, prevention and treatment. Nature Biotechnology. 2008;26:901–08.

33. Mendelsohn NJ, Messinger YH, Rosenberg AS, Kishnani PS. Elimination of antibodies to recombinant enzyme in Pompe’s disease. NEJM. 2009;360;194–5.

34. Hunley TE, Corzo D, Dudek M, Kishnani P, Amalfitano A, Chen YT, Richards SM, Phillips JA, Fogo AB, Tiller GE. Nephrotic syndrome complicating D-glucosidase replacement therapy for Pompe disease. Ped. 2004;114:e5325.

35. Banugaria SG, Prater N, Patel TT, DeArmey SM, Milleson C, Sheets KB, Bali DS, Rehder CW, Raiman JA, Wang RA, Labarthe F, Charrow J, Harmatz P, Chakraborty P, Rosenberg AS, Kishnani PS. Algorithm for the early diagnosis and treatment of patients with cross reactive immunologic material-negative classic Infantile Pompe disease: A step towards improving the efficacy of ERT. PLoS ONE. 2013;8:e67052–12.

36. Nayak S, Doerfler PA, Porvasnik SL, Cloutier DD, Khanna R, Valenzano KJ, Herzog RW, Byrne BJ. Immune responses and hypercoagulation in ERT for Pompe disease are mutation and rhGAA dose dependent. PLoS ONE. 2014;9:e98336–47.

37. de Vries JM, Kuperus E, Hoogeveen-Westerveld M, Kroos MA, Wens CAA, Stok M, van der Beek NAME, Kruijshaar ME, Rizopoulos D, van Doorn PA, van der Ploeg AT, Pim Pijnappel WWM. Pompe disease in adulthood: effects of antibody formation on enzyme replacement therapy. Genet Med. 2016;19:90–97.

38. Ohashi T, Iizuka S, Shimada Y, Eto Y, Ida H, Hachimura S, Kobayashi H. Oral administration of recombinant human acid α-glucosidase reduces specific antibody formation against enzyme in mouse. Molecular Genetics and Metabolism. 2011;103:98**–**100.

39. Parini R, De Lorenzo P, Dardis A, Burlina A, Cassio A, Cavarzere P, Concolino D, Della Casa R, Deodato F, Donati MA, Fiumara A, Gasperini S, Menni F, Pagliardini V, Sacchini M, Spada M, Taurisano R, Grazia Valsecchi M, Di Rocco M, Bembi B. Long term clinical history of an Italian cohort of infantile onset Pompe disease treated with enzyme replacement therapy. Orphanet Journal of Rare Diseases. 2018;13:32–44.

40. Nayak S, Doerfler PA, Porvasnik SL, Cloutier DD, Khanna R, Valenzano KJ, Herzog RW, Byrne BJ. Immune responses and hypercoagulation in ERT for Pompe disease are mutation and rhGAA dose dependent. PLoS ONE. 2014;9:e98336–47.

41. Diaz-Manera J, Kishnani PS, Kushlaf H, Ladha S, Mozaffar T, Straub V, Toscano A, van der Ploeg AT, Berger KI, Clemens PR, Chien Y-H, Day JW, Illarioshkin S, Roberts M, Attarian S, Borges JL, Bouhour F, Choi YC, Erdem-Ozdamar S, Goker-Alpan O, Kostera-Pruszczyk A, Haack KA, Hug C, Huynh-Ba O, Johnson J, Thibault N, Zhou T, Dimachkie MM, Schoser B, on behalf of the COMET Investigator Group. Safety and efficacy of avalglucosidase alfa versus alglucosidase alfa in patients with late-onset Pompe disease (COMET): a phase 3, randomised, multicentre trial. Lancet Neurol. 2021;20:1012–26.

42. Zhu Y, Li X, Kyazike J, Zhou Q, Thurberg BL, Raben N, Mattaliano RJ, Cheng SH. Conjugation of mannose 6-phosphate-containing oligosaccharides to acid α-glucosidase improves the clearance of glycogen in Pompe mice. J Biological Chemistry. 2004;279:50336–50341.

43. Zhu Y, Jiang JL, Gumlaw NK, Zhang J, Bercury SD, Ziegler RJ, Lee K, Kudo M, Canfield WM, Edmunds T, Jiang C, Mattaliano RJ, Cheng SH. Glycoengineered acid α-glucosidase with improved efficacy at correcting the metabolic aberrations and motor function deficits in a mouse model of Pompe disease. Mol Ther. 2009;17:954–63.

44. Zhou Q, Stefano JE, Harrahy J, Finn P, Avila L, Kyazike J, Wei R, van Patten SM, Gotschall R, Zheng X, Zhu Y, Edmunds T, Clark Q. Pan Strategies for neoglycan conjugation to human acid α-glucosidase. Bioconjug Chem. 2011;22:741–51.

45. Cohen JL, Chakraborty P, Fung-Kee-Fung K, Schwab ME, Bali D, Young SP, Gelb MH, Khaledi H, DiBattista A, Smallshaw S, Moretti F, Wong D, Lacroix C, El Demellawy D, Strickland KC, Lougheed J, Moon-Grady A, Lianoglou BR, Harmatz P, Kishnani PS, MacKenzie TC. *In utero* enzyme-replacement therapy for infantile-onset Pompe’s disease. N Engl J Med. November 11, 2022;DOI: 10.1056/NEJMoa2200587.

46. van der Ploeg AT. Science behind the study-prenatal enzyme-replacement therapy. N Eng J Med. November 11, 2022;1–5.

47. Gutschmidt K, Musumeci O, Díaz-Manera J, Chien Y-H, Knop KC, Wenninger S, Montagnese F, Pugliese A, Tavilla G, Alonso-Pérez J, Wuh-Liang Hwu P, Toscano A, Schoser B. STIG study: real-world data of long-term outcomes of adults with Pompe disease under enzyme replacement therapy with alglucosidase alfa. J Neurol. 2021; 5:1–11.

48. Korlimarlaa A, Spiridigliozzib GA, Stefanescua M, Austina SL, Kishnania PS. Behavioral, social and school functioning in children with Pompe disease. Molecular Genetics and Metabolism Reports. 2020;25:100635–12.

49. Raben N, Jatkar T, Lee A, Lu N, Dwivedi S, Nagaraju K, Plotz PH. Glycogen stored in skeletal but not in cardiac muscle in acid α-glucosidase mutant (Pompe) mice is highly resistant to transgene-encoded human enzyme. Molecular Therapy. 2002;6:601–608.

50. Neufeld EF. Lysosomal storage diseases. Annu Rev Biochem. 1991;60:257–280.

51. van den Hout JM, Reuser AJ, de Klerk JB, Arts WF, Smeitink JA, van der Ploeg AT. Enzyme therapy for Pompe disease with recombinant human α-glucosidase from rabbit milk. J Inherit Metab Dis. 2001;24:266–274.

52. Reuser AJ, Kroos MA, Hermans MM, Bijvoet AG, Verbeet MP, Van Diggelen OP, Kleijer WJ, van der Ploeg AT. Glycogenosis type II (acid maltase deficiency). Muscle Nerve. 1995;3S:61–69.

53. van den Hout H, Reuser AJ, Vulto AG, Loonen MC, Cromme-Dijkhuis A, van der Ploeg AT. Recombinant human α-glucosidase from rabbit milk in Pompe patients. Lancet. 2000;356:397–398.

54. Amalfitano A, Bengur AR, Morse RP, Majure JM, Case LE, Veerling DL, Mackey J, Kishnani P, Smith W, McVie-Wylie A, Sullivan JA, Hoganson GE, Phillips JA 3rd, Schaefer GB, Charrow J, Ware RE, Bossen EH, Chen YT. Recombinant human acid α-glucosidase enzyme therapy for infantile glycogen storage disease type II: results of a phase I/II clinical trial. Genet Med. 2001;3:132–138.

55. Corti M, Liberati C, Smith BK, Lawson LA, Tuna IS, Conlon TJ, Coleman KE, Islam S, Herzog RW, Fuller DD, Collins SW, Byrne BJ. Safety of intradiaphragmatic delivery of adeno-associated virus mediated alpha-glucosidase (rAAV1-CMV-hGAA) gene therapy in children affected by Pompe disease. Hum Gene Ther Clin Dev. 2017;28:208–1.

56. Miyamoto Y, Etoh Y, Joh R, Noda K, Ohya I, Morimatsu M. Adult-onset acid maltase deficiency in siblings. Acta Pathol Jpn. 1985;35:1533–42.

57. Anneser JMH, Pongratz DE, Podskarbi T, Shin YS, Schoser BGH. Mutations in the acid α-glucosidase gene (M. Pompe) in a patient with an unusual phenotype. Neurology. 2005;64:368–70.

58. Refai D, Lev R, Cross DT, Shimony JS, Leonard JR. Thrombotic complications of a basilar artery aneurysm in a young adult with Pompe disease. Surg Neurol. 2008;70:518–20.

59. Kretzschmar HA, Wagner H, Hubner G, Danek A, Witt TN, Mehraein P. Aneurysms and vacuolar degeneration of cerebral arteries in late-onset acid maltase deficiency. J Neurol Sci. 1990;98:169–83.

60. Makos MM, McComb RD, Hart MN, Bennett DR. Alpha-glucosidase deficiency and basilar artery aneurysm: report of a sibship. Ann Neurol. 1987;22:629–33.

61. Kobayashi H, Shimada Y, Ikegami M, Kawai T, Sakurai K, Urashima T, Ijima M, Fujiwara M, Kaneshiro E, Ohashi T, Eto Y, Ishigaki K, Osawa M, Kyosen SO, Ida H. Prognostic factors for the late onset Pompe disease with enzyme replacement therapy: from our experience of 4 cases including an autopsy case. Mol Genet Metab. 2010;100:14–9.

62. McCall AL, Dhindsa JS, Bailey AM, Pucci LA, Strickland LM, Eimallah MK. Glycogen accumulation in smooth muscle of a Pompe disease mouse model. J Smooth Muscle Res. 2021;57:8–18.

63. Hundsberger T, Rohrbach M, Kern L, Rösler KM. Swiss national guideline for reimbursement of enzyme replacement therapy in late-onset Pompe disease. J Neurol. 2013;260:2279–85.

64. Martiniuk F, Reggi S, Tchou-Wong K-M, Rom WN, Busconi M, Fogher C. Production of a functional human acid maltase in tobacco seeds: biochemical analysis, uptake by human GSDII cells, and *in vivo* studies in GAA knockout mice. Appl Biochem Biotechnol. 2013;171:916–926.

65. Martiniuk F, Mack A, Martiniuk J, Miller S, Voronin GO, Reimer D, Rossi N, Sheppard Bird L, Saleh S, Gupta R, Nigro M, Meinke P, Schoser B, Wu F, Kambitsis A, Arvanitopoulos J, Arvanitopoulos E, Tchou-Wong K-M. Preclinical studies for plant-based oral enzyme replacement therapy (Oral-ERT) in Pompe disease knockout mice with transgenic tobacco seeds expressing human GAA (tobrhGAA). https://biorxiv.org/cgi/content/short/2021.11.11.468227v1, 2021.

65a. Tisdale A, Cutillo CM, Nathan R, Russo P, Laraway B, Haende M, Nowak D, Hasche C, Chan C-H, Griese E, Dawkins H, Shukla O, Pearce DA, Rutter JL, Pariser AR. The IDeaS initiative: pilot study to assess the impact of rare diseases on patients and healthcare systems. Orphanet J Rare Dis. 2021;16:429–447.

65b. Bick D, Bick S, Fowler TA, Caulfield MJ, Scott RH. An online compendium of treatable genetic disorders, Am J Med Gen. 2020;197:48–54.

65c. Owen MJ, Wright MS, Batalov S, Kwon Y, Ding Y, Chau KK, Chowdhury S, Sweeney NM, Kiernan E, Richardson A, Batton E, Baer RJ, Bandoli G, Gleeson JG, Bainbridge M, Chambers CD, Kingsmore SF. Reclassification of the etiology of infant mortality with whole-genome sequencing. JAMA Network Open. 2023;6:e2254069.

66. Tognacca R, Botto JF. Post-transcriptional regulation of seed dormancy and germination: Current understanding and future directions. Plant Communications. 2021;2:1–22.

67. Hardy J. General aspects and applications of gibberelins and gibberellic acid in plants. In: Hardy, J., Gibberellins and Gibberellic Acid:Biosynthesis, Regulation and Physiological Effects. ed. Hauppauge: Nova Science Publishers, 2015;1–21.

68. Yu S, Wang J-W. The Crosstalk between microRNAs and gibberellin signalling in plants. Plant and Cell Physiology. 26 August 2020.

69. Briggs DE, Biochemistry of barley germination action of gibberellic acid on barley endosperm. Brewing Industry Research Foundation. 1963;69:13–19.

70. Rodrigues Vieira CA, Das Gracas Guimaraes Carvalho Vieira M, Fraga AC, Almir Oliveira J, Dos Santos CD. Action of gibberellic acid (GA3) on dormancy and activity of α-amylase in rice seeds. Revista Brasileira de Sementes. 2002;24:43–48.

71. Chen SSC, Park WM. Early actions of gibberellic acid on the embryo and on the endosperm of *Avena fatua* seeds. Plant Physiol. 1973;52:174–176.

72. Briggs DE, Effects of gibberellic acid on barley germination and its use in malting: a review. Brewing Industry Research Foundation. 1963;69:244–248.

73. Riley JM. Gibberellic acid for fruit set and seed germination. CRFG J. 1987;19:10–12.

74. Tibbot BK, Henson CA, Skadsen RW. Expression of enzymatically active, recombinant barley α-glucosidase in yeast and immunological detection of α-glucosidase from seed tissue. Plant Molecular Biology. 1998;38:379–391.

75. Frandsen TP, Lok F, Mirgorodskaya E, Roepstorff P, Svensson B. Purification, enzymatic characterization, and nucleotide sequence of a high-isoelectric-point α-glucosidase from barley malt. Plant Physiology. 2000;123:275–286.

76. Muslin EH, Kanikula AM, Clark SE, Henson CA. Overexpression, purification, and characterization of a barley α-glucosidase secreted by *Pichia pastoris*. Protein Expression and Purification. 2000;18:20–26.

77. Andriotis VME, Saalbach G, Waugh R, Field RA, Smith AM. The maltase involved in starch metabolism in barley endosperm Is encoded by a single gene. PLoS ONE. 2016;11:e0151642.

78. 2016 Exercise.com.

79. Shaaltiel Y, Bartfeld D, Hashmueli S, Baum G, Brill-Almon E, Galili Orly Dym G, Boldin-Adamsky SA, Silman I, Sussman JL, Futerman AH, Aviezer D. Production of glucocerebrosidase with terminal mannose glycans for enzyme replacement therapy of Gaucher’s disease using a plant cell system. Plant Biotechnology J. 2007;5:579–590.

80. Dressman JB, Amidon GL, Reppas C, Shah VP. Dissolution testing as a prognostic tool for oral drug absorption:immediate release dosage forms. Pharm Res. 1998;15:11–22.

81. Martiniuk F, Hirschhorn R. Characterization of neutral isozymes of human alpha-glucosidase. Biochim Biophys Acta. 1981;658:248–261.

82. Martiniuk F, Honig J, Hirschhorn R. Further studies of the structure of human placental acid alpha-glucosidase. Arch Biochem Biophys. 1984;231:454–460.

83. DeBarsy T, Jacquemin P, Devos P, Hers HG. Rodent and human acid alpha glucosidase. Purification, properties and inhibition by antibodies. Investigation in type II glycogenosis. Eur J Biochem. 1065;31:156–165.

84. Kakkis E, Lester T, Yang R, Tanaka C, Anand V, Lemontt J, Peinovich M, Passage M. Successful induction of immune tolerance to enzyme replacement therapy in canine mucopolysaccharidosis I. Proc Natl Acad Sci. 2004;101:829–834.

85. White RR, Crawley FE, Vellini M, Rovati LA. Bioavailability of 125I bromelain after oral administration to rats. Biopharm Drug Dispos. 1988;9:397–403.

86. van Hove JK, Yang HW, Wu J-Y, Brady RO, Chen YT. High-level production of recombinant human lysosomal acid α-glucosidase in Chinese hamster ovary cells which targets to heart muscle and corrects glycogen accumulation in fibroblasts from patients with Pompe disease. Proc Natl Acad Sci. 1996;93:65–70.

87. Fuller M, van der Ploeg A, Reuser AJJ, Anson DS, Hopwood JJ. Isolation and characterisation of a recombinant, precursor form of lysosomal acid α-glucosidase. Eur J Biochem. 1995;234:903–900.

88. Hirschhorn R, Reuser AJJ. Glycogen storage disease type 2 acid alpha-glucosidase acid maltase deficiency. Macgraw-Hill, Chapter 135, 2001;3389–3420.

89. Chadalavada DM, Sivakami S. Purification and biochemical characterisation of human placental acid α-glucosidase. Biochemistry and Molecular Biology International. 1997;42:1051–1061.

90. Bijvoet AGA, Kroos MA, Pieper FR, van der Vliet M, De Boer HA, van der Ploeg AT, Verbeet M, Reuser AJJ. Recombinant human acid α-glucosidase: high level production in mouse milk, biochemical characteristics, correction of enzyme deficiency in GSDII KO mice. Human Molecular Genetics. 1998;7:1815–1824.

91. van Diggelen OP, Oemardien LF, van der Beek NAME, Kroos MA, Wind HK, Voznyi YV, Burke D, Jackson M, Winchester BG, Reuser AJJ. Enzyme analysis for Pompe disease in leukocytes; superior results with natural substrate compared with artificial substrates. J Inherit Metab Dis. 2009;32:416–423.

92. Okumiya T, Keulemans JLM, Kroos MA, van der Beek NME, Boer MA, Takeuchi H, van Diggelen OP, Reuser AJJ. A new diagnostic assay for glycogen storage disease type II in mixed leukocytes. Molecular Genetics and Metabolism. 2006;88:22–28.

93. Porto C, Ferrara MC, Meli M, Acampora E, Avolio V, Rosa M, Cobucci-Ponzano B, Colombo G, Moracci M, Andria G, Parenti G. Pharmacological enhancement of α-glucosidase by the allosteric chaperone N-acetylcysteine. Molecular Therapy. 2012;20:2201–2211.

94. Khanna R, Flanagan JJ, Feng J, Soska R, Frascella M, Pellegrino LJ, Lun Y, Guillen D, Lockhart DJ, Valenzano KJ. The pharmacological chaperone AT2220 increases recombinant human acid α-glucosidase uptake and glycogen reduction in a mouse model of Pompe disease. PLoS ONE. 2012;7:e40776.

95. Martiniuk F, Chen A, Mack A, Donnabella V, Slonim A, Bulone L, Arvanitopoulos E, Raben N, Plotz P, Rom WN. Helios gene gun particle delivery for therapy of acid maltase deficiency. DNA and Cell Biol. 2002;21:717–725.

96. Martiniuk F, Bodkin M, Tzall S, Hirschhorn R. Identification of the base pair responsible for a α glucosidase allele with lower affinity for glycogen (GAA 2) and transient gene expression in deficient cells. Am J Hum Genet. 1990;47:440–445.

97. Martiniuk F, Ellenbogen A, Hirschhorn K, Hirschhorn R. Further regional localization of the genes for human acid alpha glucosidase (GAA), peptidase D (PEPD) and alpha mannosidase (MANB) by somatic cell hybridization. Hum Genet. 1985;69:109–111.

98. Mellquist JL, Kasturi L, Spitalnik SL, Shakin-Eshleman SH. The amino acid following an Asn-X-Ser/Thr sequon is an important determinant of N-linked core glycosylation efficiency. Biochemistry. 1998;37:6833–6837.

99. Moreland RJ, Jin X, Zhang XK, Decker RW, Albee KL, Lee KL, Cauthron RD, Brewer K, Edmunds T, Canfield WM. Lysosomal acid α-glucosidase consists of four different peptides processed from a single chain precursor. JBC. 2005;280:6780–6791.

100. Moreland RJ, Higgins S, Zhou A, van Straten P, Cauthron RD, Brem M, McLarty BJ, Kudo M, Canfield WM. Species-specific differences in the processing of acid α-glucosidase are due to the amino acid identity at position 201. Gene. 2012;491:25–30.

101. Martiniuk F, Chen A, Donnabella V, Arvanitopoulos E, Slonim AE, Raben N, Plotz P, Rom WN. Correction of glycogen storage disease type II by enzyme replacement with a recombinant human acid maltase produced by over-expression in a CHO-DHFR (neg) cell line. Biochem Biophys Res Commun. 2000;276:917–923.

102. Hintze S, Limmer S, Dabrowska-Schlepp P, Berg B, Krieghoff N, Busch A, Schaaf A, Meinke P, Schoser B. Moss-derived human recombinant GAA provides an optimized enzyme uptake in differentiated human muscle cells of Pompe disease. Int J Mol Sci. 2020;21:2642–2657.

103. Raben N, Nagaraju K, Lee E, Kessler P, Byrne B, Lee L, LaMaurex M, King J, Sauer B, Plotz P. Targeted disruption of the acid alpha glucosidase gene in mice causes an illness with critical features of both infantile and adult human glycogen storage disease type II. JBC. 1998;273:19086–92.

104. Wheeler TL, Eppolito AK, Smith LN, Huff TB, Smith RF. A novel method for oral stimulant administration in the neonate rat and similar species. J Neuroscience Methods. 2007;159:282–285.

105. Gomez-Lechon MJ, Ponsoda X, Castell JV. A microassay for measuring glycogen in 96-well-cultured cells. Analytical Biochemistry. 1996;236:296–301.

106. Bian G-L, Wei L-C, Shi M, Wang YQ, Cao R, Chen LW. Fluoro-Jade C can specifically stain the degenerative neurons in the substantia nigra of the 1-methyl-4-phenyl-1,2,3,6-tetrahydro pyridine-treated

107. Maga JA, Zhou J, Kambampati R, Peng S, Wang X, Bohnsack RN, Thomm A, Golata S, Tom P, Dahms NM, Byrne BJ, LeBowitz JH. Glycosylation-independent lysosomal targeting of acid α-glucosidase enhances muscle glycogen clearance in pompe mice. J Biol Chem. 2013;288:1428–38.

108. Zhu Y, Li X, McVie-Wylie A, Jiang C, Thurberg BL, Raben N, Mattaliano RJ, Cheng SH. Carbohydrate-remodelled acid α-glucosidase with higher affinity for the cation-independent mannose 6-phosphate receptor demonstrates improved delivery to muscles of Pompe mice. Biochem J. 2005;389:619–628.

109. McVie-Wylie AJ, Lee KL, Qiu H, Jin X, Do H, Gotschall R, Thurberg BL, Rogers C, Raben N, O’Callaghan M, Canfield W, Andrews L, McPherson JM, Mattaliano RJ. Biochemical and pharmacological characterization of different recombinant acid α-glucosidase preparations evaluated for the treatment of Pompe disease. Mol Genet Metab. 2008;94:448–455.

110. Tong PY, Kornfeld S. Ligand interactions of the cation-dependent mannose 6-phosphate receptor. Comparison with the cation-independent mannose 6-phosphate receptor J Biol Chem. 1989;264:7970–7975.

111. Fukuda T, Ewan L, Bauer M, Mattaliano RJ, Zaal K, Ralston E, Plotz PH, Raben N. Dysfunction of endocytic and autophagic pathways in a lysosomal storage disease. Ann Neurol. 2006;59:700–708.

112. Raben N, Schreiner C, Baum R, Takikita S, Xu S, Xie T, Myerowitz R, Komatsu M, van Der Meulen J, Nagaraju K, Ralston E, Plotz P. Suppression of autophagy permits successful enzyme replacement therapy in a lysosomal storage disorder-murine Pompe disease. Autophagy. 2010;6:1078–1089.

113. Lim J-A, Li L, Shirihai OS, Trudeau KM, Rosa Puertollano R, Raben N. Modulation of mTOR signaling as a strategy for the treatment of Pompe disease. EMBO Molecular Medicine. 2017;9:353–370.

114. Carroll B, Maetzel D, Maddocks OD, Otten G, Ratcliff M, Smith GR, Dunlop EA, Passos JF, Davies OR, Jaenisch R, Tee AR, Sarkar S and Korolchuk VI. Control of TSC2-Rheb signaling axis by arginine regulates mTORC1 activity. eLife. 2016;5:e11058–91.

115. Jung JW, Kim NS, Jang SH, Shin YJ, Yang MS. Production and characterization of recombinant human acid alpha-glucosidase in transgenic rice cell suspension culture. J Biotechnol. 2016;226:44**–**53.

116. Jung J-W, Huy N-X, Kim H-B, Kim N-S, Giap DV, Moon-Sik Yang. Production of recombinant human acid α-glucosidase with high-mannose glycans in gnt1 rice for the treatment of Pompe disease. J Biotechnology. 2017;249:42–50.

117. Sariyatun R, Florence J, Kajiura H, Ohashi T, Misaki R, Fujiyama K. Production of human acid-alpha glucosidase with a paucimannose structure by glycoengineered *Arabidopsis* cell culture. Frontiers in Plant Science. 2021;12:1–13.

118. Cheung SCK, Liu L, Lan L, Liu Q, Sun SSM, Chan JCN, Tong PCY. Glucose lowering effect of transgenic human insulin-like growth factor-I from rice: *in vitro* and *in vivo* studies. BMC Biotech. 2011;11:37–47.

119. Hashizume F, Hino S, Kakehashi M, Okajima S, Nadano D, Aoki N, Matsuda T. Development and evaluation of transgenic rice seeds accumulating a type II-collagen tolerogenic peptide. Transgenic Res. 2008;17:1117–1129.

120. Keum YS, Khor TO, Lin W, Shen G, Kwon K H, Barve A, Li A, Koug AN. Pharmacokinetics and pharmacodynamics of broccoli sprouts on the suppression of prostate cancer in transgenic adenocarcinoma of mouse prostate (TRAMP) mice: implication of induction of Nrf2, H0-1 and apoptosis and the suppression of Akt-dependent kinase pathway. Pharma Res. 2009;26:2324–2331.

121. Lamphear BJ, Streatfield SJ, Jilka JM, Brooks CA, Barker DK, Turner DD, Delaney DE, Garcia M, Wiggins B, Woodard SL, Hooda EE, Tizard IR, Lawhorn B, Howard JA. Delivery of subunit vaccines in maize seed. J Contr Release. 2002;85:169–180.

122. Ning T, Xie T, Qiu Q, Yang W, Zhou S, Zhou L, Zheng C, Zhu Y, Yang D. Oral administration of recombinant human granulocyte macrophage colony stimulating factor expressed in rice endosperm can increase leukocytes in mice. Biotech Lett. 2008;30:1679–1686.

123. Pena Ramırez YJ, Tasciotti E, Gutierrez-Ortega A, Donayre Torres AJ, Olivera Flores MT, Giacca M, Gomez Lim MA. Fruit-specific expression of the human immunodeficiency virus type 1 Tat gene in tomato plants and Its immunogenic potential in mice. Clin Vaccine Immunol. 2007;14:685–692.

124. Sardana RK, Alli Z, Dudani A, Tackaberry E, Panahi M, Narayanan M, Ganz P, Altosaar I. Biological activity of human granulocyte-macrophage colony stimulating factor is maintained in a fusion with seed glutelin peptide. Transgenic Res. 2002;11:521–531.

125. Streatfield SJ, Lane JR, Brooks CA, Barker DK, Poage ML, Mayor JM, Lamphear BJ, Drees CF, Jilka JM, Hood EE, Howard JA. Corn as a production system for human and animal vaccines. Vaccine. 2003;21:812–815.

126. Suzuki K, Kaminuma O, Yang L, Takai T, Mori A, Umezu-Goto M, Ohtomo M, Ohmachi Y, Noda Y, Hirose S, Okumura K, Ogawa H, Takada K, Hirasawa M, Hiroi T, Takaiwa F. Prevention of allergic asthma by vaccination with transgenic rice seed expressing mite allergen: induction of allergen-specific oral tolerance without bystander suppression. Plant Biotechnology Journal. 2011;9:982–990.

127. Tackaberry ES, Dudani AK, Prior F, Tocchi M, Sardana R, Altosaar I, Ganz PR. Development of biopharmaceuticals in plant expression systems: cloning, expression and immunological reactivity of human cytomegalovirus glycoprotein B (UL55) in seeds of transgenic tobacco. Vaccine. 1990;17:3020–3029.

128. Takagi H, Hiroi T, Yang I, Tada Y, Yuki Y, Takamura K, Ishimitsu R, Kawauchi H, Kiyono H, Takaiwa F. A rice-based edible vaccine expressing multiple T cell epitopes induces oral tolerance for inhibition of Th2-mediated IgE responses. Proc Natl Acad Sci. 2005;102:17525–17530.

129. Takagi H, Hiroi T, Yang L, Takamura K, Ishimitsu R, Kawauchi H, Takaiwa F. Efficient induction of oral tolerance by fusing cholera toxin B subunit with allergen-specific T-cell epitopes accumulated in rice seed. Vaccine. 2008;26:6027–6030.

130. Takagia H, Hiroi T, Hirosea S, Yanga L, Takaiwaa F. Rice seed ER-derived protein body as an efficient delivery vehicle for oral tolerogenic peptides. Peptides. 2009;31:1421–1425.

131. Takaiwa F. Update on the use of transgenic rice seeds in oral immunotherapy. Immunotherapy. 2013;5:301–12.

132. Wu J, Yu L, Li L, Hu J, Zhou J, Zhou X. Oral immunization with transgenic rice seeds expressing VP2 protein of infectious bursal disease virus induces protective immune responses in chickens. Plant Biotech J. 2007;5:570–578.

133. Xie T, Qiu Q, Zhang W, Tingting Ning T, Yang W, Zheng C, Wanga C, Zhu Y, Yang D. A biologically active rhIGF-1 fusion accumulated in transgenic rice seeds can reduce blood glucose in diabetic mice via oral delivery. Peptides. 2008;29:1862–1870.

134. Yamada Y, Nishizawa K, Yokoo M, Zhao H, Onishi K, Teraishi M, Utsumi S, Ishimoto M, Yoshikawa M. Anti-hypertensive activity of genetically modified soybean seeds accumulating novokinin. Peptides. 2008;29:331–337.

135. Yang L, Kajiura H, Suzuki H, Hirose S, Fujiyama K, Takaiwa F. Generation of a transgenic rice seed-based edible vaccine against house dust mite allergy. Bioch Biophys Res Comm. 2008;365:334–339.

136. Yang L, Tada Y, Yamamoto MP, Zhao H, Yoshikawa M, Takaiw F. A transgenic rice seed accumulating an anti-hypertensive peptide reduces the blood pressure of spontaneously hypertensive rats. FEBS Lett. 2006;580:3315–3320.

137. Yang L, Wakasa Y, Takaiwa F. Biopharming to increase bioactive peptides in rice seed. JAOAC International. 2008;91:957–66.

138. Zimmermann J, Saalbach I, Jahn D, Giersberg M, Haehnel, S, Wedel J, Macek J, Zoufal K, Glunder G, Falkenburg D, Kipriyanov SM. Antibody expressing pea seeds as fodder for prevention of gastrointestinal parasitic infections in chickens. BMC Biotech. 2009;9:79–101.

139. Kaufman J. The economic potential of plant-made pharmaceuticals in the manufacture of biologic pharmaceuticals. J Commercial Biotechnology. 2011;17:173–182.

140. Grabowski GA, Golembo M, Shaaltiel Y. Taliglucerase alfa: an enzyme replacement therapy using plant cell expression technology. Mol Gen Met. 2014;112:1–8.

141. Tekoah Y, Shulman A, Kizhner T, Ruderfer I, Fux L, Nataf Y, Bartfeld D, Ariel T, Gingis**–**Velitski S, Hanania U, Shaaltiel Y. Large-scale production of pharmaceutical proteins in plant cell culture**—**the Protalix experience. Plant Biotechnology J. 2015;13:1199–1208.

142. Shaaltiel Y, Gingis**–**Velitski S, Tzaban S, Fiks N, Tekoah Y, Aviezer D. Plant-based oral delivery of β-glucocerebrosidase as an enzyme replacement therapy for Gaucher’s disease. Plant Biotechnology J. 2015;13:1033–1040.

143. Paul M, Ma J. Plant-made pharmaceuticals:leading products and production. Biotech Applied Biochem. 2011;58:58–67.

144. Kwon K-C, Daniell H. Low-cost oral delivery of protein drugs bioencapsulated in plant cells. Plant Biotechnol J. 2015;13:1017–1022.

